# Efficient DNA base editing via an optimized DYW-like deaminase

**DOI:** 10.1101/2024.05.15.594452

**Authors:** Jiyeon Kweon, Soomin Park, Mi Yeon Jeon, Kayeong Lim, Gayoung Jang, An-Hee Jang, Minyoung Lee, Cheong Seok, Chaeyeon Lee, Subin Park, Jiseong Ahn, JiYoon Jang, Young Hoon Sung, Daesik Kim, Yongsub Kim

**Author notes:** Correspondence (J.K.), (D.K.), (Y.K.). These authors contributed equally (Jiyeon Kweon, Soomin Park).

## Abstract

CRISPR-based cytosine base editors enable precise genome editing without inducing double-stranded DNA breaks, yet traditionally depend on a limited selection of deaminases from the APOBEC/AID or TadA families. Here, we introduce SsCBE, a novel CRISPR-based cytosine base editor utilizing SsdA_tox_, a DYW-like deaminase derived from the toxin of *Pseudomonas syringae*. Strategic engineering of SsdA_tox_ has led to remarkable improvements in the base editing efficiency (by up to 8.4-fold) and specificity for SsCBE, while concurrently reducing cytotoxicity. Exhibiting exceptional versatility, SsCBE was delivered and efficiently applied using diverse delivery methods, including the engineered virus-like particles (eVLPs). Its application has enabled targeted cytosine base editing in mouse zygotes and pioneering edits in mitochondrial DNA. The advent of SsCBE marks a significant advancement in the CRISPR toolkit, providing a versatile tool for advanced research and therapeutic strategies.

## INTRODUCTION

Base editing, a cutting-edge development in the field of genome engineering, represents a significant leap beyond traditional genome editing tools. It allows for the precise and targeted alteration of nucleotide sequences without the need for DNA double-stranded breaks or donor templates(1). Central to the mechanism of base editors (BEs) are its two main components: DNA-binding modules, such as the CRISPR-Cas system, TALEs, and ZFPs, and the deaminase enzymes that facilitate direct nucleotide conversion. CRISPR-based base editing exemplifies this evolution, employing a fusion of Cas9 with single-stranded DNA (ssDNA) targeting deaminases, such as those from the AID/APOBEC family or engineered TadA variants(2-4). The Cas9-gRNA complex binds the targeted DNA sequence and forms an R-loop, exposing ssDNA. The exposed strand is then edited by the deaminase, converting specific nucleotides with high precision. An alternative strategy incorporates TALEs or ZFPs with dsDNA-targeting deaminases like DddAtox derived from the SCP1.201 family, further diversifying the base editing toolkit and expanding its potential applications for organellar base editing(5-8).

Despite these advancements, the range of deaminases currently employed in base editing is somewhat limited. Predominantly, members of the AID/APOBEC family are utilized in Cytosine Base Editors (CBEs), TadA variants are employed in both CBEs and Adenine Base Editors (ABEs), and DddA variants are used in double-stranded DNA-targeting CBEs (DsCBEs)(9,10) (11) (12,13). In this study, we aim to further expand the toolkit of base editing by introducing a novel, compact, and highly efficient deaminase from the DYW-like deaminase, integrated with Cas9 and TALE systems. We fused nCas9 with SsdA_tox_, known as a bacterial toxin derived from *Pseudomonas syringae*, enabling targeted cytosine base editing and named SsCBE (SsdA_tox_-derived CBE)(14). Through rational engineering of SsdA_tox_, we successfully improved the base editing efficiency of SsCBE comparable to conventional CBE, BE4max, with reduced genome- and transcriptome-wide off-target effects. Finally, we demonstrate that SsCBE, when fused with TALE proteins, can be utilized for efficient cytosine base editing in mitochondrial DNA. Our findings contribute to expanding the base editing toolkit, paving the way for innovative applications in genome editing.

## Methods

### Plasmid construction

The human codon optimized SsdA_tox_ domain was synthetized (Integrated DNA Technologies) and cloned into either the N-terminus or C-terminus of a modified pCMV_BE4max vector (Addgene #112093) with the UGI domain deleted. To construct SsdA_tox_–UGI variants, one or two copies of the UGI domain were amplified by Phusion High-Fidelity DNA Polymerase (Thermo Fisher Scientific) and cloned into designated positions. For constructing rationally engineered SsCBE-UGI-C2 variants, primers containing mismatches with wild–type sequences were used to introduce mutations into the SsdA_tox_ sequences. The amplicons were then cloned into the wild-type SsCBE-UGI-C2 vector using Gibson Assembly Master Mix (New England Biolabs). The cjSsCBE2 was constructed by exchanging APOBEC1 domain of cjCBEmax to SsdA_tox_-SRE domain, and the pAAV-cjABE8e-gRNA-ANGPT2-HPD-2 was modified to construct the single AAV vector encoding jSsCBE2(15). The gRNAs were constructed using pRG2Z vector (Addgene #104174) and pU6-Cj-sgRNA (Addgene #89753).

### Cell culture and transfection

HEK293T/17 (ATCC CRL–11268) and HeLa (ATCC CCL–2) were maintained in DMEM medium with 10% fetal bovine serum (FBS) and 1% penicillin–streptomycin. K562 (ATCC CCL–243) and SKOV3 (ATCC HTB–77) were maintained in RPMI and McCoy’s 5A medium, respectively, with 10% FBS and 1% penicillin–streptomycin. All the mammalian cells were incubated at 37 °C in a 0.05% CO_2_ atmosphere and routinely tested for Mycoplasma contamination using MicoStrip (InvivoGen). The cells were seeded onto 48–well plate (Corning) one day before transfection, and transfection was conducted at 50–60% cell confluency using Lipofectamine 2000 (Thermo Fisher Scientific) unless otherwise stated.

Briefly, a total of 500 ng plasmid DNA (250 ng of gRNA and 250 ng of base editors or each 250ng of TALE-SREs) were mixed with 1.5 μL Lipofectamine 2000. For K562 cells, 2×105 cells were electroporated with 250 ng of gRNA plasmids and 750 ng of CBE using the SF Cell Line Nucleofector X Kit (Lonza) via the 4D-Nucleofector system. Genomic DNA was extracted 96h after transfection using homemade cell lysis buffer (10 mM Tris-HCl pH8.5 and 0.05% SDS) or DNeasy Blood & Tissue Kits (Qiagen) to evaluate editing frequencies. For evaluating cell viability, cells 72h after transfection were subjected to luminescent assay using CellTiter-Glo 2.0 (Promega) according to the manufacturer’s protocol. To compare the expression levels of each SsdA_tox_ variants, the P2A-mcherry fused variants were transfected and subjected to FACS (BD FACSCanto) analysis 48h after transfection.

### Targeted deep sequencing

Genomic DNA containing either the on-target or off-target sites was amplified with KAPA HiFi HotStart DNA polymerase (Roche) or SUN-PCR Blent (SUN GENETICS) according to the manufacturer’s instructions. The amplified products, including Illumina TruSeq HT dual index adapter sequences, were subjected to 150 bp paired-end sequencing using the Illumina iSeq 100 platform. MAUND, an analysis tool accessible at https://github.com/ibs-cge/maund, was used to determine the base editing efficiencies.

### AAV particle production and transduction

HEK293T/17 cells were seeded onto 150 mm culture dish one day before transfection, and pAAV-cjSsCBE2-ANGPT2 or pAAV-cjSsCBE2-HPD was transfected with pAAV-DJ and helper plasmids. The transfection was conducted at 70% cell confluency and with plasmids at the molar ratio of 1:1:1. AAV particles were collected and concentrated 72h after transfection using AAVpro Purification Kit Midi (Takara) according to manufacturer’s protocol. 1×10^4^ HEK293T/17 cells were seeded in 96-well plate and AAV particles at different vg/cell were transduced. Cells were collected 96h after transduction to measure cytosine base editing frequency. The vg/cell was determined by real-time PCR using AAVpro Titration Kit (Takara) according to the manufacturer’s protocol. HEK293T/17 cells using in the experiments regarding AAV production and transduction were maintained in DMEM with 2% FBS.

### Protein purification

The plasmid encoding the pABE8e-protein (Addgene plasmid #161788) was used to construct the plasmid encoding the His_8_-SsCBE2-C2, by Gibson assembly method. The transformed Rosetta cells (EMD Millipore) were grown overnight at 37 °C in Luria-Bertani (LB) broth supplemented with 100 μg/ml kanamycin and 34 μg/ml chloramphenicol after transformation with the His_8_-SsCBE2-C2 plasmid DNA. Subsequently, 10 milliliters of overnight cultures of Rosetta cells transformed with His_8_-SsCBE2-C2 plasmid DNA were inoculated into 400 milliliters of LB broth supplemented with 100 μg/ml kanamycin and 34 μg/ml chloramphenicol at 30°C until the OD600 reached 0.5-0.6. The cells were cooled to 18°C for one hour, followed by induction of His_8_-SsCBE2-C2 protein with 0.8% rhamnose, and subsequent culture for another 18 hours. For protein purification, cells were harvested by centrifugation at 5,000g for 10 minutes at 4 °C and lysed via sonication in 5 ml lysis buffer (50 mM NaH_2_PO_4_, 300 mM NaCl, 1 mM DTT, and 10 mM imidazole, pH 8.0) supplemented with lysozyme (Sigma) and protease inhibitor (Roche complete, EDTA-free). The soluble lysate obtained after centrifugation at 13,000 rpm for 30 minutes at 4°C was incubated with Ni-NTA agarose resin (Qiagen) for 1 hour at 4°C. The mixture of lysate and Ni-NTA was applied onto a column and washed with a buffer containing 50 mM NaH2PO4, 300 mM NaCl, and 20 mM imidazole at pH 8.0. The SsCBE2-C2 protein was subsequently extracted by utilizing the elution buffer (50 mM NaH_2_PO_4_, 300 mM NaCl, and 250 mM imidazole, pH 8.0). To improve the purity of the SsCBE2-C2 protein, we subjected fractions containing the protein to additional purification steps. These fractions were combined with Heparin beads (Cytiva) in a solution composed of 20 mM Tris-HCl (pH 8.0 at 25 °C), 150 mM NaCl, 5% (v/v) glycerol, and 1% (v/v) Triton. We eluted proteins by a linear gradient of NaCl concentration from 600 mM to 2 M in a buffer containing 20 mM Tris-HCl (pH 8.0 at 25°C), 0.1 mM DTT, and 5% (v/v) glycerol. The fractions that contained the SsCBE2-C2 protein were collected and concentrated through an Amicon Ultra centrifugal filter (Millipore). Following that, a buffer exchange with storage buffer (20 mM HEPES-KOH, pH 7.5, 150 mM KCl, 1 mM DTT, and 20% glycerol) was performed. The concentrated protein underwent further processing with centrifugal filter units (Millipore). The purified SsCBE2-C2 protein was then stored at -80°C.

### Digenome sequencing

The genomic DNA was extracted using the DNeasy Tissue Kit (Qiagen) following the manufacturer’s protocols. To induce in vitro deamination, the SsCBE2-C2 protein (100 nM) was pre-incubated with gRNA (300 nM) for 10 minutes at room temperature. Subsequently, the preassembled complex was mixed with genomic DNA (10 μg) in a reaction buffer (100 mM NaCl, 50 mM Tris-HCl, 10 mM MgCl_2_, and 100 μg/ml BSA) to a final reaction volume of 1000 μL. The reaction mixture was incubated at 37°C for 8 hr. After deamination, the genomic DNA was purified using the DNeasy tissue kit (Qiagen), and RNase A (50 μg/mL) was added for the elimination of gRNA. A second incubation step with USER enzyme (6 units) was performed on purified genomic DNA (2 μg), and the reaction volume was 100 μL, incubated at 37°C for 3 hours. This was followed by another purification round using the DNeasy Blood & Tissue kit (Qiagen). After digestion with SsCBE2-C2 and USER enzymes, the genomic DNA was subjected to whole-genome sequencing (WGS) at 30-40x depth using an Illumina HiSeq X Ten sequencer at Macrogen. The genome sequence was mapped using the Isaac aligner, and the DNA cleavage sites were identified using the Digenome program, available at https://github.com/chizksh/digenome-toolkit2.

### Orthogonal R-loop assay

dsaCas9 (Addgene plasmid #138162) and their gRNAs were used in orthogonal R-loop assay (16). A total of 500 ng plasmids (each 100 ng plasmids of dsaCas9, sa-gRNAs, SsCBE or BE4max, gRNAs, and p3s-EFS-puromycin^R^) were transfected in HEK293T/17 cells and puromycin was treated 24h after transfection at 1μg/mL to select transfected cells. Genomic DNA was extracted 96h after transfection and editing frequencies were measured by targeted-deep sequencing.

### Transcriptome sequencing

Total RNA was extracted 48 hours after transfection using the RNeasy Mini Kit (Qiagen) according to the manufacturer’s instructions. RNA libraries were then generated with the TruSeq Stranded Total RNA Library Prep Gold kit (Illumina). Evaluation of RNA library quality was performed using the Agilent 2200 TapeStation with a D1000 ScreenTape system. Total RNA sequencing was conducted at Macrogen using a NovaSeq 6000 Sequencer (Illumina) with paired-end sequencing (2 x 100bp).

### RNA variant calling

To analyze RNA sequencing data generated by next-generation sequencing (NGS), we used a previously validated RNA variant calling pipeline designed for the analysis of off-target RNA base editing (17,18).The NGS data were aligned to the hg38 (release v.105) human reference genome using the STAR aligner (v.2.7.10a). BAM files were then processed for RNA variant calling using MarkDuplicates, BaseRecalibrator, ApplyBQSR, and HaplotypeCaller from the GATK package (v.4.2.4.1). We filtered RNA variant loci by comparing them with control samples. In experimental set of replicate #1, the untreated replicate #2 served as the control, while in experimental sets of replicate #2, the untreated replicate #1 served as the control. RNA variant loci with a variant count of at least 2 and a read depth of at least 10 were retained. We excluded RNA variant loci that were already present in the control sample or that were considered indeterminate due to low sequencing depth in the control sample. C-to-T editing was quantified as RNA variant loci with C-to-T editing on the positive strand or G-to-A editing on the negative strand among total RNA editing. Similarly, A-to-G editing was quantified as RNA variant loci with A-to-G editing on the positive strand or T-to-C editing on the negative strand among total RNA editing.

### SsCBE2-C2 eVLP production and purification

The plasmid containing pCMV-MMLVgag-3xNES-ABE8e was a gift from David Liu (Addgene plasmid #181751). Construction of the plasmid encoding pCMV-MMLVgag-3xNES-SsCBE2-C2 was accomplished through Gibson assembly. SsCBE2-C2-eVLPs were generated by transfecting Gesicle Producer 293T cells. Gesicle cells were seeded in a 150 mm cell culture dish at a density of 5×10^6^ cells per dish. After 20–24 hours, the cells were transfected using PEI (sigma) according to the manufacturer’s protocols. For producing SsCBE2-C2 eVLP, a mixture of plasmids expressing VSV-G (400 ng), MMLVgag–pro–pol (3,375 ng), MMLVgag–3xNES–SsCBE2-C2-eVLPs (1,125 ng), and a gRNA (4,400 ng) were co-transfected per 150 mm cell culture dish. At 40–48 hours post-transfection, the supernatant from transfected cells was harvested and subjected to centrifugation for 5 minutes at 500 g to eliminate cell debris. The resulting supernatant was then filtered through a 0.45-μm PVDF filter. For concentration, the filtered supernatant underwent a 100-fold concentration step using PEG-it Virus Precipitation Solution (System Biosciences), according to the manufacturer’s protocols.

### Construction of TALE-SsCBE

Target specific TALE-SsCBEs were assembled by one step golden gate cloning with TALE RVD modules(19). Empty expression vectors were first constructed according to the previous study by modifying TALEN expression vectors(5). Briefly, fokI domains were exchanged by SsdA_tox_-SRE domain to construct TALE-SsCBE expression vectors. For nucleus genome targeting TALE-SsCBEs, two copies of UGI were cloned between the NLS and N-terminal domains of TALE (NTD). For mitochondrial genome targeting TALE-SsCBE, MTS was cloned instead of NLS and one copy of UGI and NES signal were added in C-terminus of expression vectors. All the assembled TALE-SsCBEs were subjected to Sanger sequencing to confirm their sequences.

### Generation of *Tyr*^*Q68\**^ mutant mice

All animal experiments were conducted according to the Korean Ministry of Food and Drug Safety (MFDS) guidelines, and animal protocols were reviewed and approved by the Institutional Animal Care and Use Committees (IACUC) of Asan Institute for Life Sciences (permit number: 2020-14-347). All mice were maintained in the specific pathogen-free (SPF) facility of the Laboratory of Animal Research in the Asan Medical Center (AMCLAR).

To construct the *in vitro* transcription template for the gRNA, the following pair of oligomers was annealed and cloned into pUC57-sgRNA vector (Addgene plasmids #51132): 5′−TAGGACCTCAGTTC CCCTTCAAAG−3′ and 5′−AAAC CTTTGAAGGGGAACTGAGGT−3′. The constructs encoding BE3 or SsCBE2-C2 were linearized using the PmeI restriction enzyme (NEW ENGLAND BioLabs). mRNAs of base editors and gRNA were synthesized *in vitro* from linearized DNA templates using the mMESSAGE mMACHINE T7 Ultra kit (Ambion) and MEGAshortscript T7 kit (Ambion), respectively, according to the manufacture protocol.

C57BL/6N (DBL, Republic of Korea) and ICR (OrientBio, Republic of Korea) mouse strains were used as embryo donors and foster mothers, respectively. Micromanipulation of fertilized eggs collected from female mice of C57BL/6N (B6N) strain and subsequent processes required for the mouse model establishment were performed as previously described(20). The mixture of BE3 or SsCBE2-C2 mRNA (250 ng/μl) and gRNA (50 ng/μl) was diluted with RNase-free injection buffer (0.25 mM EDTA, 10 mM Tris, pH 7.4) and microinjected into male pronuclei using a TransferMan NK2 micromanipulator and a FemtoJet microinjector (Eppendorf). The manipulated embryos were transferred into the oviducts of pseudopregnant foster mothers. Genomic DNA samples were isolated from tail biopsies of newborn mice or E13.5 mouse embryos were analyzed by targeted-deep sequencing.

### Statistical analysis

All results are expressed as the mean ± SEM unless indicated otherwise. Statistical analysis was performed in GraphPad Prism 9.1.1. p values were derived using Student’s two-tailed t testing.

## RESULTS

### Evaluating a novel toxin–derived deaminase

We focused on a newly identified interbacterial cytidine deaminase toxin from the DYW-like subgroup encoded by *P. syringae*, named SsdA_tox_(21) (Figure 1A). Mongous and co-workers demonstrated that SsdA_tox_ can induce C:G to T:A transitions in E. coli, and showed in vitro deaminating activity toward single-stranded DNA (ssDNA)(14). The SsdA_tox_ comprises 151 amino acid (aa) residues, making it approximately 66% shorter than the rAPOBEC1 deaminase domain (229 aa) used in representative CBEs, BE3 and BE4max(2,22). We initially confirmed SsdA_tox_’s divergence from other deaminases used in CRISPR-mediated base editing tools through phylogenetic tree analysis (Supplemental Figure S1). To validate the deaminase activity of the SsdA_tox_ domain, we purified it and conducted an *in vitro* deamination assay. Using a FAM-labeled ssDNA substrate, we confirmed that SsdA_tox_ domain has cytosine–to–uracil conversion activity (Supplemental Figure S2). We then incubated SsdA_tox_ protein with genomic DNA substrate from HEK293T/17 cells, dCas9 protein, and gRNA, and analyzed the nucleotide sequences of target sites using targeted deep sequencing. The dCas9 and gRNA form an R-loop at genomic DNA target sites, exposing ssDNA substrates for SsdA_tox_-induced cytosine-to-uracil conversion. We evaluated cytosine–to–thymine conversion frequencies at two target sites, RNF2 and HEK2, since uracil is read as thymine in sequencing. Targeted deep sequencing revealed that in the HEK2 and RNF2 sequences, despite the activity of SsdA_tox_ is restricted to the non-target strand and limiting conversion to 50%, cytosines were converted to thymine with a maximum frequency of 18.5% (Figure 1B and Supplemental Figure S3A). Targeted deep sequencing revealed that these conversion frequencies to thymine decreased to background levels after treatment with the uracil-Specific Excision Reagent (USER) enzyme, confirming that SsdA_tox_ protein indeed induced cytosine-to-uracil conversion, as the USER enzyme can recognize and eliminate uracil. These results underscore the potential of the SsdA_tox_ domain for use in cytosine base editors.

**Figure 1.**
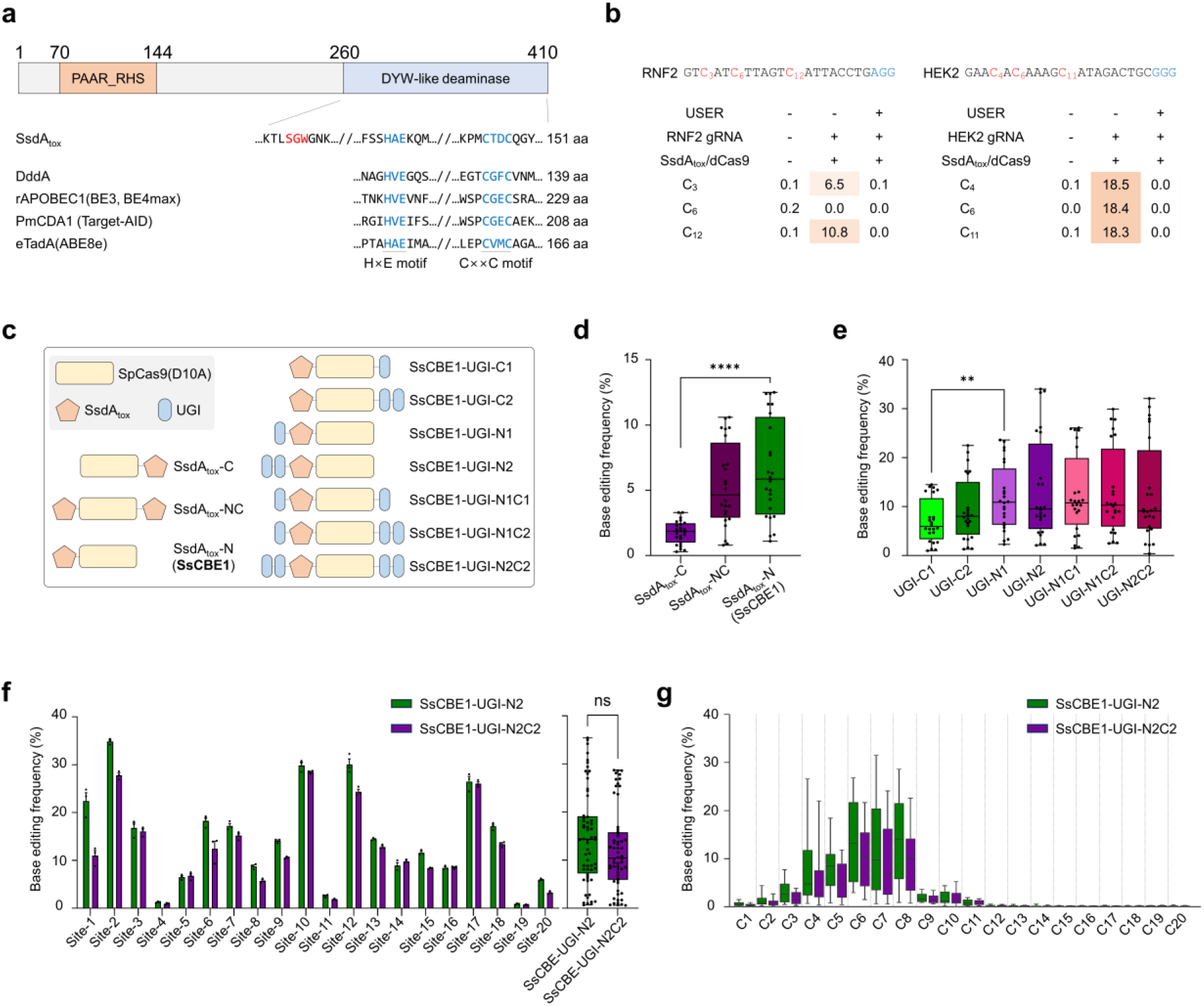
Use of the interbacterial toxin SsdA_tox_ from the DYW-like deaminase clade in CRISPR cytosine base editor toolkits. **a** Domain analysis of full-length SSdA_tox_, highlighting PAAR (proline-alanine-alanine-arginine), RHS (recombination hotspots), and the DYW-like deaminase toxin domain, now referred to as SSdA_tox_ in this study. **b** *in vitro* deaminase assay utilizing the SSdA_tox_ domain showed C to T conversion efficiencies at the RNF2 ((C3, C6, and C12) and HEK2 (C4, C6, and C11) target sites measured by targeted deep sequencing. The maximum conversion efficiency reached up to 50%, limited to editing of cytosine in the target strand only. **c** Schematic overview of SSdA_tox_ -based cytosine base editors. The constructs combine SpCas9 (D10A) nickase, SsdAtox domain, and UGI domains. **d, e** Base editing frequencies of each construct across 8 target sites were depicted in box-whisker plot. Dots represent the independent biological triplicate of each 8 target sites. ** P=0.0056, ***** P<0.0001 by unpaired t-test. **f** Base editing frequencies of SsCBE1-UGI-N2 and SsCBE1-UGI-N2C2 in HEK293T/17 UNG KO cells. Data are presented as mean and error bars indicate SEM of independent biological triplicate in bar graph (n=3). Base editing frequencies were compared in box-whisker plot and unpaired *t*-test was used to statistical analysis. **g** Base editing window analysis of both constructs across 19 target sites.

### Development of a toxin–based novel cytosine base editor

To develop a novel cytosine base editor, we constructed several combinations of SsdA_tox_, uracil-DNA glycosylase inhibitor (UGI), and spCas9-D10A nickase domains (Figure 1C). First, we evaluated the base editing frequencies of three constructs, SsdA_tox_-C (SsdA_tox_ fused to the C-terminal of spCas9-D10A nickase), SsdA_tox_-N (SsdA_tox_ fused to the N-terminal of spCas9-D10A nickase), and SsdA_tox_-NC (SsdA_tox_ fused to both the N- and C-terminal of spCas9-D10A nickase), across eight endogenous sites in *UNG* knockout HEK293T/17 cells (Figure 1D). We found that SsdA_tox_-C exhibited relatively lower cytosine base editing activity compared to the other two constructs, with SsdA_tox_-N reaching up to 12.4% cytosine base editing frequency at the HEK3 target site (Supplemental Figure S3B). To further optimize, we added the UGI domain to either the N-terminus or C-terminus of the SsdA_tox_-N domain (hereafter referred to as SsCBE1) and examined their base editing efficiency across eight endogenous sites in *UNG* knockout HEK293T/17 cells (Figure 1E and Supplemental Figure S3C). Although the base editing frequencies of UGI variants were examined in *UNG* knockout backgrounds, we observed that UGI composition slightly affected base editing frequencies; SsCBE1 with N-terminal UGI domains showed higher base editing frequencies than those with C-terminal UGI domains. These results suggest that UGI composition might affect protein expression of the construct itself or other factors involved in the base editing mechanism, thereby altering base editing frequencies. To confirm the versatility of the SsCBE1-based CBE, we chose UGI-N2 (two UGIs fused to the N-terminal of SsCBE1) and UGI-N2C2 (the two UGIs fused to both the N- and C-terminal of SsCBE1) variants and evaluated the base editing frequencies across an additional 19 endogenous sites (Figure 1F). The UGI-N2 variant showed up to 35.4% cytosine base editing frequency at the Site-2 target site, with both UGI-N2 and UGI-N2C2 variants displaying similar base editing frequencies across the 20 target sites. We further analyzed the base editing windows of the two SsCBE1 variants across 28 endogenous sites in HEK293T/17 cells (Figure 1G). We confirmed that both variants exhibited high base editing frequencies at cytosine positions from 4 to 8 (numbered 1–20 in the 5′ to 3′ direction),, similar to the base editing window of the representative rAPOBEC1-based CBE(2).

To further evaluate the characteristics of SsCBE1-UGI-N2 (hereafter referred to as SsCBE1-N2), we transfected the SsCBE1-N2 construct into wild-type HEK293T/17 cells and measured the base editing frequencies across 6 endogenous sites (Figure 2A). The SsCBE1-N2 exhibited base editing frequencies up to 24.2% at the HEK2 target site; however, the base editing frequencies of SsCBE1-N2 were lower than those of the representative CBE, BE3 and BE4max. (Supplemental Figure S4A). We also evaluated the product purity of SsCBE1-N2 and confirmed that it generally had high product purity, though there was room for improvement; at the cytosine position 6 of HEK2 target sequences, SsCBE-N2 had better product purity than BE3 but not than BE4max (Supplemental Figure S4B). Therefore, we decided to engineer the SsdA_tox_ domain to develop cytosine base editors with improved performance.

**Figure 2.**
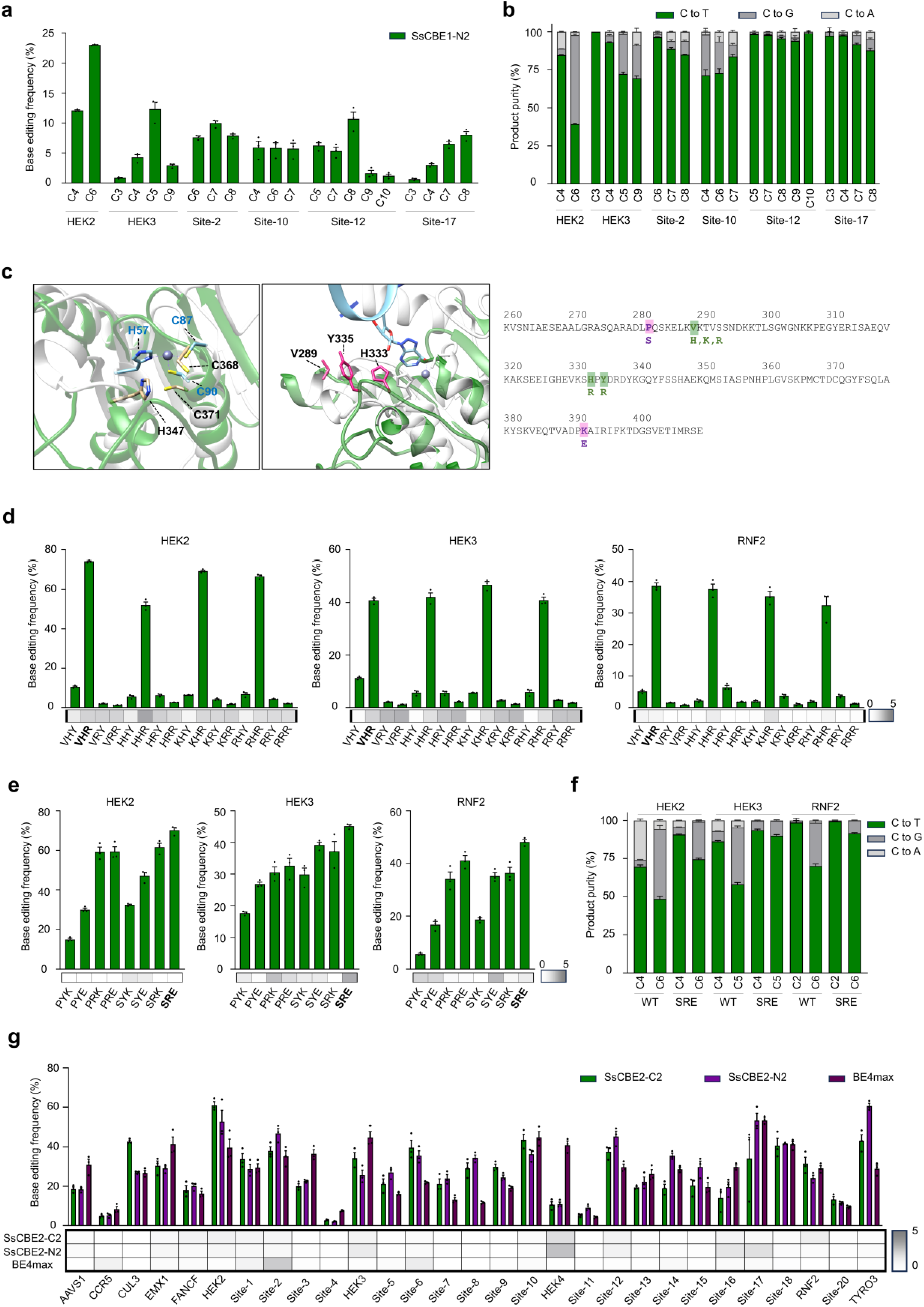
Enhancement of cytosine base editing activity in HEK293T/17 cells through rational engineering of the SsdA_tox_ domain. **a** Base editing frequency of SsCBE1-N2 in HEK293T/17 cells. Data are presented as mean and error bars indicate SEM of independent biological triplicate (n=3). **b** Product purity of SsCBE1-N2 at each cytosine position. **c** Structural alignment of TadA from ABE8e (gray, PDB ID:6VPC) and SsdA_tox_ (green, PDB ID: 7JTU). Left panel: Catalytic active sites (Blue for TadA, black for SsdA_tox_). Middle panel: Three candidate resides for engineering (V289, H333, and Y335). Right panel: Five candidate positions for engineering were highlighted and designed amino acid residues were listed below. **d-e** Base editing and indel frequencies of engineered variants; **d**, rational engineering at V289, H333 and Y335 positions; **e**, and P282 and K392. Data are presented as mean and error bars mean SEM of independent biological triplicate samples (n=3). Heatmaps below each bar plot showed indel frequencies. **f** Comparison of product purity between WT and SRE variant across three target sites. **g** Base editing frequencies of SsCBE2-C2, SsCBE2-N2, and BE4max across 29 endogenous target sites in HEK293T/17 cells. Data are presented as mean and error bar means SEM of independent biological triplicate samples (n=3). Indel frequencies were described below each bar plot.

### Rational engineering of SsdA_tox_

We compared the structure of the SsdA_tox_ protein (Protein Data Bank [PDB] ID: 7JTU) with that of ABE8e (PDB ID: 6VPC), which contains a similarly sized and well-characterized TadA deaminase (TadA8e), to predict their active sites and DNA-binding moieties(14,23) (Figure 2C). The active sites of SsdA_tox_ resemble those of TadA8e, and based on the merged structure, three residues (V289, H333, Y335) were identified for potential enhancement of substrate interaction.

We initially cloned five variants (V289H, V289K, V289R, H333R, and Y335R) and ten combined variants of SsdA_tox_ into the SsCBE1-UGI-C2 construct (hereafter referred to as SsCBE1-C2) and evaluated their base editing frequencies across three endogenous sites in HEK293T/17 cells (Figure 2D). Remarkably, the Y335R mutation increased cytosine base editing frequencies at the HEK2, HEK3, and RNF2 sites by 7.0-, 3.6-, and 7.5-fold, respectively, compared to wild-type SsdA_tox_. Further evaluation of seven variants with combinations of P282S, K392E, and Y335R mutations showed that both P282S and K392E mutations enhanced base editing frequencies (Figure 2E and Supplemental Figure S4C). The final engineered SsdA_tox_ variant, featuring mutations P282S, Y335R, and K392E (named the SRE variant), exhibited an average 4.1-fold improvement in cytosine base editing efficiency and enhanced product purity compared to the wild-type in HEK293T/17 cells across three endogenous sites (Figure 2F and Supplemental Figure S4D). Given that the SsdA_tox_ domain is derived from a toxin, we evaluated the cytotoxicity and the expression of several constructs in HEK293T/17 cells. Interestingly, protein engineering improved both the cytotoxicity and the expression levels of SsdA_tox_ domain; the SRE variants appeared to be a similar cell viability to BE4max (Supplemental Figure S5). We then incorporated the SRE variants into the SsCBE1-N2 construct, named SsCBE-N2, and evaluated their base editing efficiency in HEK293T/17 cells across 29 endogenous target sites (Figure 2G and Supplemental Figure S4E). While wild-type SsdA_tox_ demonstrated enhanced performance with an N-terminal UGI domain, SRE variants showed comparable performance with both N-terminal and C-terminal UGI domains. Additionally, both SsCBE2-N2 and SsCBE2-C2 exhibited comparable base editing efficiency and indel frequencies to those of BE4max (Supplemental Figure S4E). The base editing window of these variants appeared to be cytosine position from 4 to 8, with slight editing observed at cytosine position 3 and 9 of target sequences (Supplemental Figure S4F). As is known, BE4max showed lower base editing frequency in a G<u>C</u> context(2); however, SsCBE2 did not exhibit a context preference (Supplemental Figure S4G). We also demonstrated that SsCBE2-C2 could induce high-frequency cytosine base editing in other cell lines, including K562, SKOV3, and HeLa cells (Supplemental Figure S6). These findings indicate that rational engineering of the SsdA_tox_ domain leads to significant improvements, yielding base editing frequencies comparable to those of BE4max.

### gRNA dependent off-target effects of SsCBE2

To assess the specificity of SsCBE2, we first investigated whether SsCBE2-C2 can tolerate mismatches in the spacer sequence of gRNAs. We transfected HEK293T/17 cells with plasmids encoding SsCBE2-C2 or BE4max, together with plasmids encoding the corresponding gRNA, each containing zero to four mismatches in the spacer sequences. We then determined the substitution frequencies at two endogenous target sites (Figure 3A and Supplemental Figure S7A). Overall, SsCBE2-C2 generally exhibited less tolerance for mismatched targets than BE4max. For instance, the relative frequencies of SsCBE2-C2 and BE4max-induced substitutions at mismatched versus matched sites were 0.05 and 0.60, respectively, for a gRNA containing a single mismatch (at position 6, numbered 1-20 in the 5’ to 3’ direction) at the RNF2 site (Supplemental Figure S7A). These results suggest that SsCBE2-C2 enables precise genome editing with a lower tolerance for mismatches compared to BE4max.

**Figure 3.**
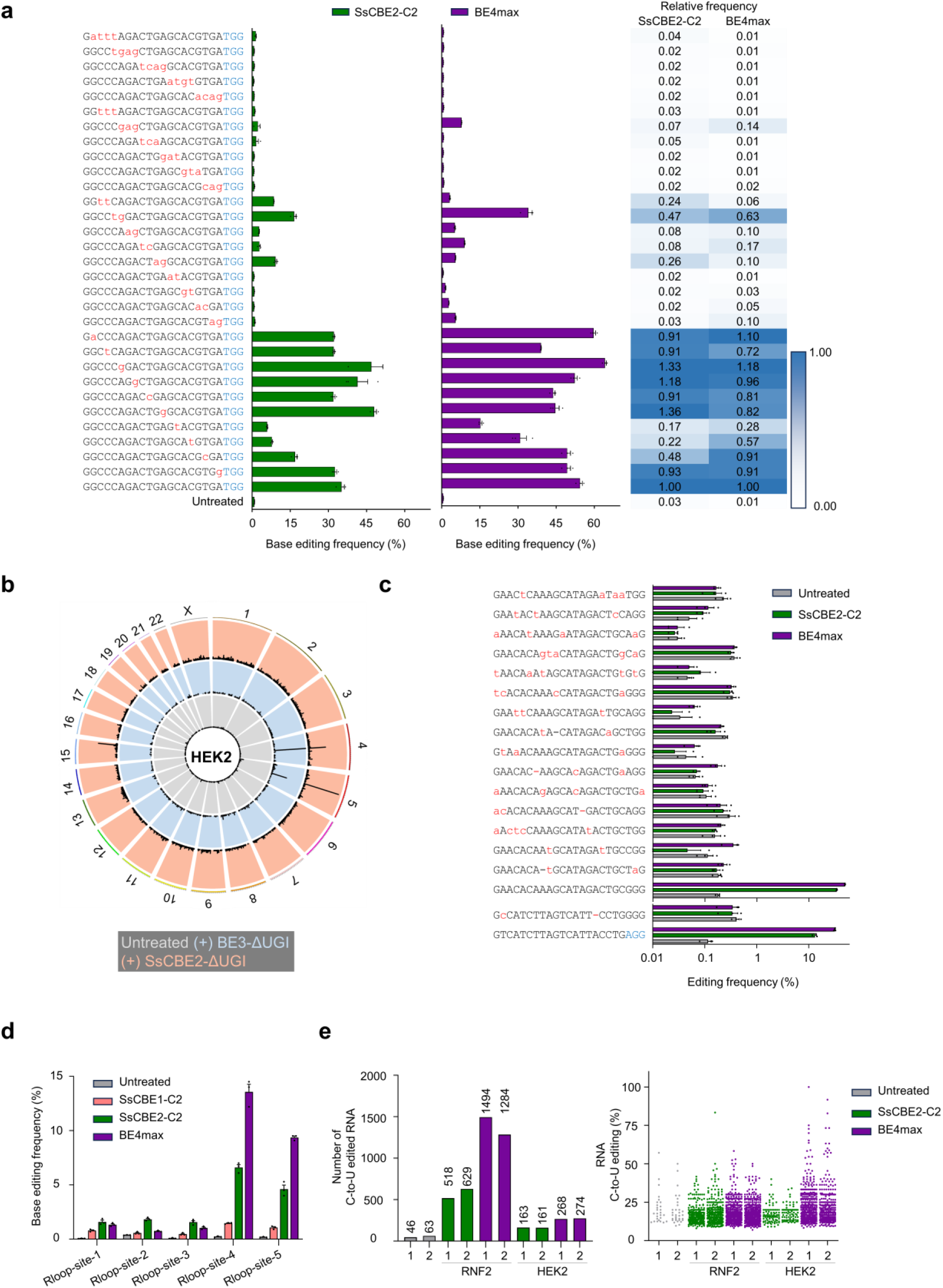
Identification of off-target effects of SsCBE2-C2. **a** Mismatch tolerance of SsCBE2-C2 and BE4max towards sgRNAs with one to four nucleotides mismatches from the HEK3 site in HEK293T/17 cells. PAM sequences are indicated in blue and mismatched bases are indicated in red. Relative frequencies were calculated by dividing base editing frequencies obtained with mismatched sgRNAs by the mean base editing frequency of the matched sgRNA. Data are presented as mean and error bar means SEM of independent biological triplicate samples (n=3). **b** Representative Circos plot illustrating genome-wide DNA cleavage scores obtained by Digenome-seq. Digenome-seq was performed with intact genomic DNA (gray), BE3-ΔUGI plus hAAG and Endo VIII (blue), or with SsCBE2-ΔUGI plus hAAG and Endo VIII (red). **c** Editing frequencies at off-target sites captured by Digenome-seq were measured by targeted-deep sequencing in HEK293T/17 cells. PAM sequences are indicated in blue and mismatched bases are indicated in red. Dashes represent RNA bulges. Data are presented as mean and error bar means SEM of independent biological triplicate (n=3). **d** Measurement of Cas9-independent off-target deamination by dSaCas9-mediated orthogonal R-loop assay in HEK293T/17 cells. Data are presented as mean and error bar represent SEM of independent biological triplicate (n=3). **e** Cas9-independent RNA off-target deamination of SsCBE2-C2 and BE4max in HEK293T/17 cells. Transcriptome sequencing was used to determine the number of C-to-U edited nucleotides and the frequency of RNA C-to-U editing.

To determine whether Digenome-seq could effectively evaluate the genome-wide target specificities of SsCBE2-C2, we incubated human genomic DNA from HEK293T/17 cells with ribonucleoproteins (RNPs) composed of purified SsCBE2-C2 protein and in vitro-transcribed gRNA, followed by treatment with the USER enzyme(24) (Supplemental Figure S7B). After whole genome sequencing the digested genomic DNA, we aligned the sequence reads to the human reference genome and used the integrative genomics viewer (IGV) to examine the alignment patterns at the on-target site(25). The alignments indicated specific cleavage by SsCBE2-C2 and USER enzymes at the on-target site (Supplemental Figure S7B). To identify off-target sites of SsCBE2-C2 in the human genome, we utilized a DNA cleavage score previously employed in our research (Figure 3B)(24,26,27). Based on the Digenome-seq method, we observed 2 and 16 potential off-target sites for SsCBE2-C2 targeting RNF2 and HEK2, respectively (Supplemental Figure S7C and Supplemental Table S1). To validate the off-target effects captured by Digenome-seq, we used targeted deep sequencing and determined SsCBE2-C2 and BE4max-induced substitution frequencies in HEK293T/17 cells. We examined 15 potential off-target sites and confirmed that none of these sites were validated by targeted deep sequencing with SsCBE2-C2 (Figure 3C). In contrast, BE4max induced two validated off-target events (HEK2-OT2 and HEK2-OT6) containing either two base pair (bp) mismatches or two bp mismatches with one bp RNA bulge, with observed frequencies of 0.35% and 0.18%, respectively (Figure 3C). These findings underscore the high specificity of SsCBE2-C2.

### gRNA independent DNA and RNA off-target effects of SsCBE

To compare the gRNA-independent DNA deamination of SsCBE2-C2 and BE4max, we conducted an orthogonal R-loop assay using catalytically inactive *Staphylococcus aureus* Cas9 (dsaCas9) and saCas9 gRNA(16). We assessed the gRNA-independent DNA deamination of SsCBE2-C2 and BE4max at artificially induced single-strand DNA sites by transfecting HEK293T/17 cells with plasmid DNA encoding eighter SsCBE2-C2 or BE4max, alongside spCas9 gRNA, dsaCas9, and saCas9 gRNA. Deamination frequencies in the R-loop formed by dsaCas9 were measured across five endogenous sites using targeted deep sequencing (Figure 3D). Notably, SsCBE2-C2 exhibited higher gRNA-independent off-target deamination at sites 1, 2, and 3 compared to BE4max, but lower gRNA-independent off-target deamination at sites 5 and 6 (Figure 3D). These results indicate that the gRNA-independent DNA deamination of SsCBE2-C2 is comparable to that of BE4max (Figure 3D).

Previous studies have shown that rAPOBEC1-based CBEs cause transcriptome-wide deamination, resulting in C-to-U conversion in a gRNA-independent manner(17,28). To assess the gRNA-independent RNA off-target effects of SsCBE2-C2 and BE4max, HEK293T/17 cells were transfected with plasmids encoding SsCBE2-C2 or BE4max along with corresponding gRNAs targeting RNF2 or HEK2. Total RNA was isolated two days post-transfection, and RNA sequencing (RNA-seq) was performed to assess transcriptome-wide RNA off-target editing; the Genome Analysis Toolkit (GATK) was used for RNA variant calling. Our analysis revealed that gRNA-independent C-to-U RNA editing induced by SsCBE2-C2 is lower than that induced by BE4max (Figure 3E and Supplemental Figure S7D). Additionally, the extent of A-to-G RNA editing was similar between BE4max-treated and untreated HEK293T/17 cells (Supplemental Figure S7E). These findings suggest that the gRNA-independent RNA off-target C-to-U deamination of SsCBE2-C2 is lower than that of BE4max.

### The versatility of SsCBE1-SRE

The cjCas9 is one of the smallest Cas9 orthologs, making it a promising tool for in vivo therapy. We have previously developed a cjCas9-based CBE, cjCBEmax, which has an N-terminal rAPOBEC1 domain and a C-terminal tandem UGI domain of the cjCas9-D8A-L58Y/D900K nickase, cjCBEmax(15). We replaced rAPOBEC1 with the SRE variant to generate cjSsCBE2 and evaluated its editing efficiency in HEK293T/17 cells across four endogenous sites. The cjSsCBE2 exhibited a 2.2-fold average improvement in cytosine base editing efficiency (Figure 4A and Supplemental Figure 8A). Subsequently, we produced AAV particles, infected HEK293T/17 cells, and found that the base editing frequencies increased in a dose-dependent manner (Figure 4B and Supplemental Figure 8B).

**Figure 4.**
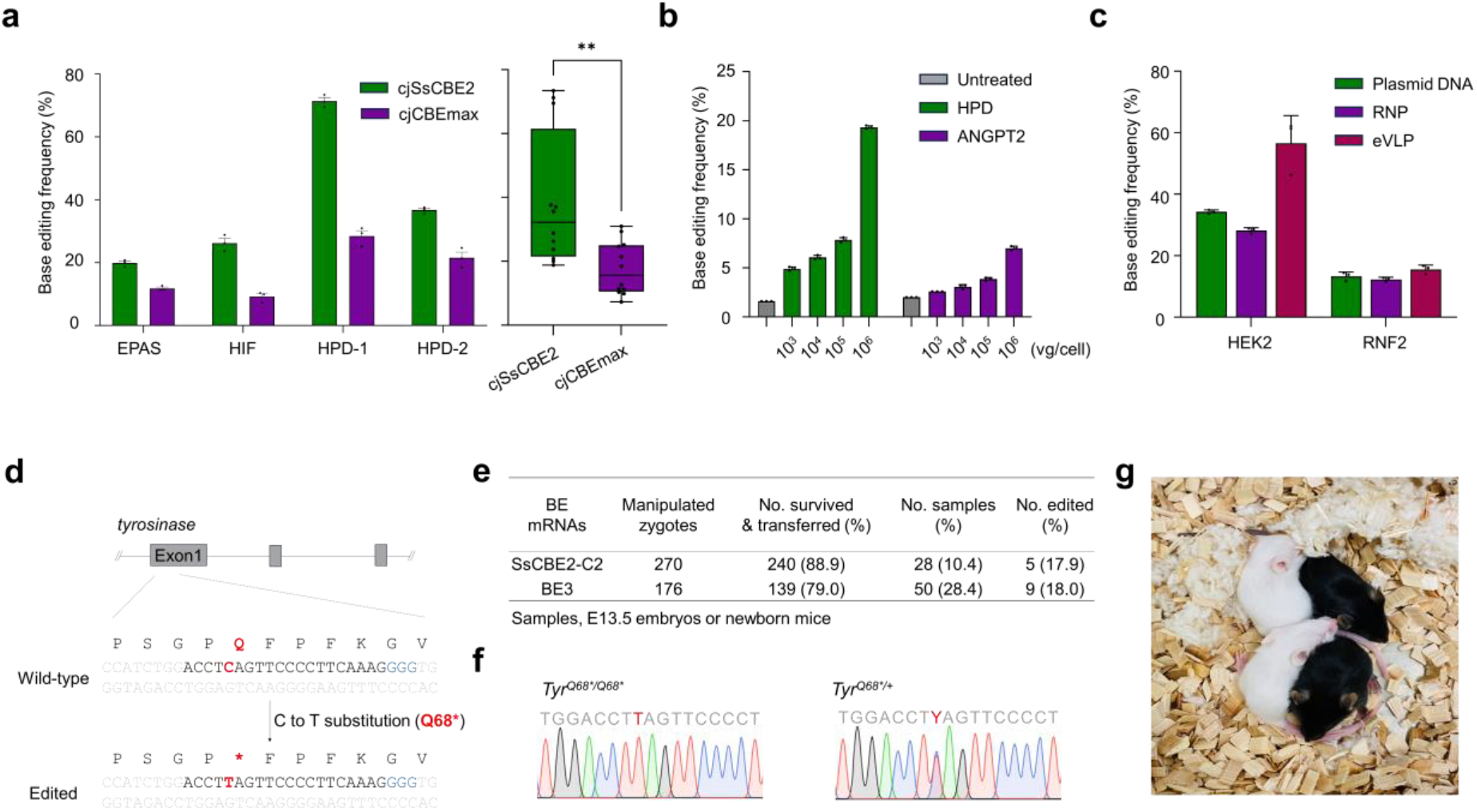
The versatility of the SsCBE2. **a** Base editing frequency of cjSsCBE2 and cjCBEmax at four target sites in HEK293T/17 cells. Data represents means and error bar indicate SEM of independent biological triplicate (n=3). ** *P*=0.0040 by unpaired *t*-test. **b** AAV particles produced with pAAV-cjSsCBE2-UGI×1-HPD or pAAV-cjSsCBE2-UGI×1-ANGPT2 were transduced at different vg/cells in HEK293T/17 cells and base editing frequencies were measured by targeted-deep sequencing. Data are plotted as mean and error bars represent SEM of independent biological triplicate (n=3). **c** Comparison of SsCBE2-C2 base editing frequencies according to three delivery methods in HEK293T/17 cells; plasmids DNA, RNP, or eVLP. Data are presented as mean and error bar means SEM of independent biological triplicate (n=3). **d** Schematic overviews of tyrosinase gene disruption by SsCBE2-mediate cytosine base editing. PAM sequences are highlighted in blue and target cytosine is indicated in red. C-to-T conversion at target cytosine induce the nonsense mutation (Q68*). **e** Summarized results of microinjection of SsCBE2-C2 or BE3 mRNA in mouse one cell embryos. **f-g** The nucleotide sequences of target sequences from pups were confirmed by Sanger sequencing. The C-to-T converted positions are highlighted in red.

Cas9 nuclease and BEs are used for RNP delivery, which is known to reduce off-target effects and cytotoxicity compared to plasmid DNA delivery(27,29,30). To enhance the specificity of SsCBE2-C2, we employed RNP delivery by transfecting preassembled SsCBE2-C2 protein and in vitro-transcribed gRNA into HEK293T/17 cells. The results indicate that SsCBE2-C2 RNP delivery showed activity similar to that of plasmid DNA delivery (Figure 4C). We then generated engineered virus-like particles (eVLP) using SsCBE2-C2 to explore in vivo gene therapy potential(31). After infecting HEK293T/17 cells with eVLPs containing SsCBE2-C2 protein and corresponding gRNA, we observed that eVLP-mediated base editing of SsCBE2-C2 exhibited comparable or higher activity than plasmid DNA delivery depending on the target sites (Figure 4C).

To investigate the cytosine base editing capabilities of SsCBE2-C2 in vivo, we tried to induce a premature stop codon through a single C-to-T conversion at the tyrosinase (*Tyr*) gene (*Tyr*^Q68*^) in mouse zygotes. As illustrated in Figure 4D, we utilized a gRNA known to specifically designed to target the mouse *Tyr* gene and co-delivered SsCBE2-C2 mRNA along with this gRNA into mouse zygotes obtained from the C57BL/6NTac (B6N) mouse strain(32). No acute toxicity was observed following the microinjection of SsCBE2-C2 mRNA into the B6 mouse zygotes (Figure 4E). We observed the edited allele in 5 out of 28 mice and embryos (17.9%), compared to a frequency of 18.0% (9 out of 50 embryos) when using the BE3, and successfully generated *Tyr*^Q68*^ mice using SsCBE2-C2 (Figure 4E-G). The developmental rate of the SsCBE2-C2 mRNA-injected mouse embryos was relatively lower than that of embryos edited with the BE3. However, given the sensitivity of B6 embryos, these findings suggest that SsCBE2-C2 is a suitable tool for gene editing in mouse fertilized eggs. This indicates that SsCBE2-C2 has potential for efficient base editing in vivo, offering an alternative to existing base editors with potentially enhanced specificity or efficiency for targeted gene modifications in mice.

### Mitochondrial DNA base editing using SRE variants

Recent biochemical studies conducted in vitro have shown that the enzyme SsdA_tox_ possesses cytosine deamination activity on ssDNA. Interestingly, at elevated concentrations, SsdA_tox_ also exhibits the ability to deaminate cytosine in dsDNA(14,33). To further explore the potential of SRE variant in mediating cytosine deamination within dsDNA genomes, we engineered fusions of SRE variant with TALE constructs, named TALE-SRE, akin to DdCBE systems, which is incorporating UGI and a nuclear localization signal (NLS)(5). These TALE arrays were assembled using the high-throughput Golden Gate assembly method(19), targeting three endogenous genomic sites (HEK3, HEK4, and TYRO3).

Upon comparing with the DdCBE system, known for inducing high-frequency base editing at targeted loci, our constructs did not exhibit significant base editing activity at these genomic sites (Supplemental Figure S9A). Subsequently, we constructed TALE arrays targeting mitochondrial DNA sites, including ND1, COX3, CYN, and ATP6, and fused these with SRE variant, UGI, and a mitochondrial transport signal. Following transfection into HEK293T/17 cells, we quantified nucleotide conversion frequencies. Remarkably, the TALE-SRE-UGI constructs achieved C to T conversion rates of up to 9.7% at the four targeted mitochondrial DNA sites (Figure 5 and Supplemental Figure S9B). Contrary to the DdCBE system, which functions as a dimer, our observations indicate that TALE-SRE can operate effectively as monomers at the target sites. These findings underscore the versatility of SRE variant, highlighting its capacity to target both ssDNA and dsDNA within the eukaryotic cellular environment.

**Figure 5.**
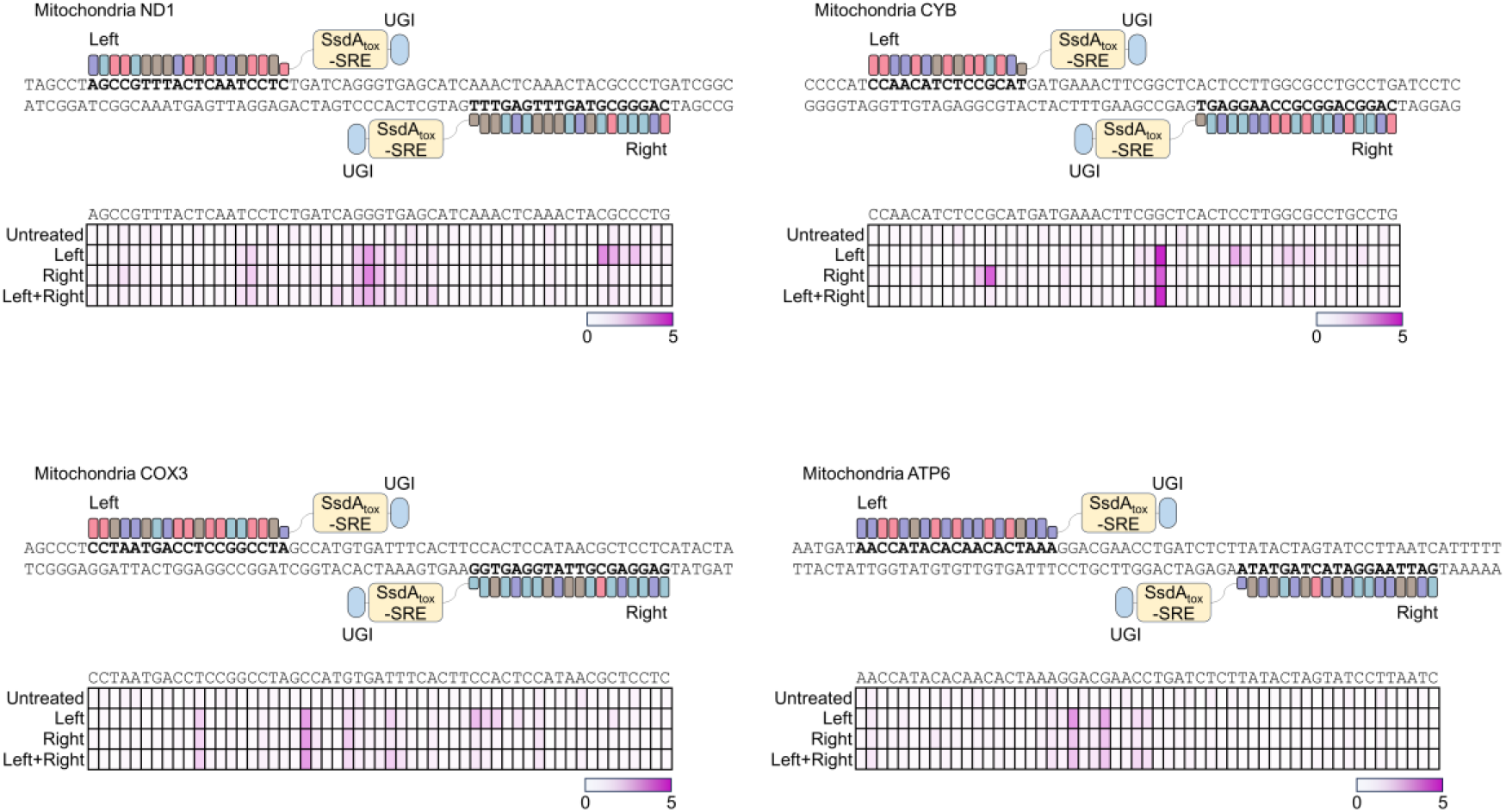
Mitochondrial genome editing using TALE-SRE in HEK293T/17 cells. Architectures of TALE-SRE targeting the ND1, CYB, COX3, and ATP6 sites were described. Monomer (left-TALE-SRE or right-TALE-SRE) and dimer (left-TALE-SRE and right-TALE-SRE) forms of TALE-SRE were transfected in HEK293T/17 cells at independent biological triplicate and base editing frequencies were measured by targeted-deep sequencing and described in heatmaps.

## Discussion

Previously identified DYW deaminase family members are known to target RNA, but, despite structural similarities to the family, SsdA_tox_ from the DYW-like deaminase subgroup shows cytosine deaminase activity toward ssDNA substrates. Here, we report the development of a novel CRISPR base editor utilizing the DYW-like deaminase. By fusing SsdA_tox_ with nCas9, we demonstrated that the SsdA_tox_-based base editor can induce cytosine base editing in human cells. Furthermore, we enhanced the cytosine base editing activity and moderated cytotoxicity through rational protein engineering of the SsdA_tox_ domain, achieving comparable activity with the current representative CBE, BE4max. Using in-depth DNA and RNA off-target analysis, we confirmed that the newly developed CBE, SsCBE2-C2, had slightly improved specificity compared to BE4max. Additionally, we showcased the versatility of the newly developed CBE by editing endogenous target sites using both AAV particles from a single cjSsCBE2 vector and DNA-free delivery methods, including RNP and eVLP delivery, highlighting the broad applicability of the SsdA_tox_-based CBE.

As SsdA_tox_ is derived from a bacterial toxin, we observed that the wild-type SsdA_tox_ domain exhibited toxicity in *E*.*coli* during sub-cloning. Although the wild-type SsdA_tox_ domain showed cytotoxicity in mammalian cells, we overcame this issue by incorporating a UGI domain and employing rational protein engineering (Supplemental Figure S5). The SsCBE2-C2 variant exhibited slightly improved performance in DNA and RNA off-target activity, suggesting that further engineering is required to minimize off-target effects.

Recently, Gao and co-workers discovered a number of small and efficient deaminases using AlphaFold2, which were utilized for cytosine base editing(34); However, the DYW-like deaminase we used, SsdA_tox_, was not included in their findings. Upon experimenting with a truncated version of SsdA_tox_, which is similar in size or slightly larger than other deaminases, we confirmed that its size could be further reduced by 5 amino acids without compromising base editing efficiency (Supplemental Figure S10).

Unexpectedly, the TALE-SRE, which may target dsDNA as a substrate, showed cytosine base editing activity, but we cannot find any genome-wide off target effect through *in vitro* digenome-sequencing. This finding is noteworthy as previous studies that dsDNA can be targetable with considerably lower efficiency compared to ssDNA by SsdA_tox_ domain. To the best of our knowledge, this is the first study to demonstrate the feasibility of nuclear and mitochondrial base editing using DYW-like deaminases highlighting the potential of a different clade of deaminases for genome editing tools. Furthermore, the development of a small CBE opens up new possibilities for base editing applications, not only in the nucleus but also in cellular organelles.

## Author contributions

Y.K supervised the research. J.K, D.K. and Y.K. conceived the research. J.K, S.P., Y.H.S., D.K. and Y.K. designed the study. J.K, S.P., D.K. and Y.K. performed and analyzed the main experiments. J.K., S.P., M.Y.J., K.L., A.-H.J., M.L., C.S., C.L., S.P, J.A., J.J. performed the experiments. J.K, S.P., Y.H.S., D.K. and Y.K. wrote the manuscript.

## Data availability

The high-throughput sequencing data from this study have been deposited in the NCBI Sequence Read Archive (SRA) database under the accession code PRJNA1076329 and is publicly available as of the date of publication.

