## Supplementary information for "Efficient DNA base editing via an optimized DYW-like deaminase"

[1] Department of Cell and Genetic Engineering, Asan Medical Institute of Convergence Science and Technology, Asan Medical Center, University of Ulsan College of Medicine, Seoul 05505, Republic of Korea
[2] Stem Cell Immunomodulation Research Center, University of Ulsan College of Medicine, Seoul 05505, Republic of Korea
[3] Department of Precision Medicine, Sungkyunkwan University School of Medicine, Suwon 16419, Republic of Korea
[4] Convergence Medicine Research Center, Asan Institute for Life Sciences, Asan Medical Center, Seoul 05505, Republic of Korea,
[5] Brain Science Institute, Korea Institute of Science and Technology (KIST), Seoul 02792, Republic of Korea
[6] Division of Bio-Medical Science & Technology, KIST School, Korea University of Science and Technology, Seoul 02792, Republic of Korea
[7] These authors contributed equally

* Correspondence:
 (J.K.), (D.K.), (Y.K.)

Table of Contents

Figure S1. Phylogenetic tree of representative 18 deaminase domains
Figure S2. In vitro deaminase assay using SsdAtox protein
Figure S3. Developments of SsdAtox-based cytosine base editors
Figure S4. Rational engineering of SsdAtox domain
Figure S5. Cytotoxicity and expression of SsdAtox variants
Figure S6. Base editing in other three cell lines
Figure S7. Analysis of off-target effects of SsCBE2
Figure S8. Architectures of cjCas9-based cytosine base editors
Figure S9. Base editing using TALE-SRE
Figure S10. Truncation of SsdAtox-SRE domain

Table S1. Identified potential off-target sites by Digenome-seq
Table S2. Nucleotide sequences of target sites and PCR primers
Table S3. Target sequences and PCR sequences for TALE-SRE

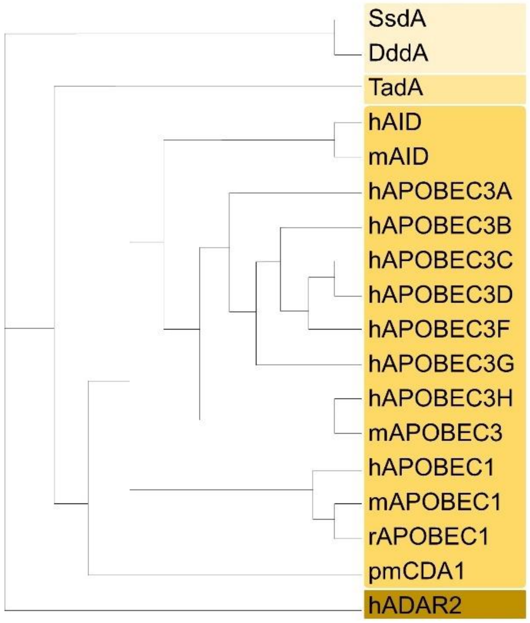


**Fig. S1. Phylogenetic tree of representative 18 deaminase domains.**
Phylogenetic tree was obtained by T-COFFEE Multiple Sequence Alignment.

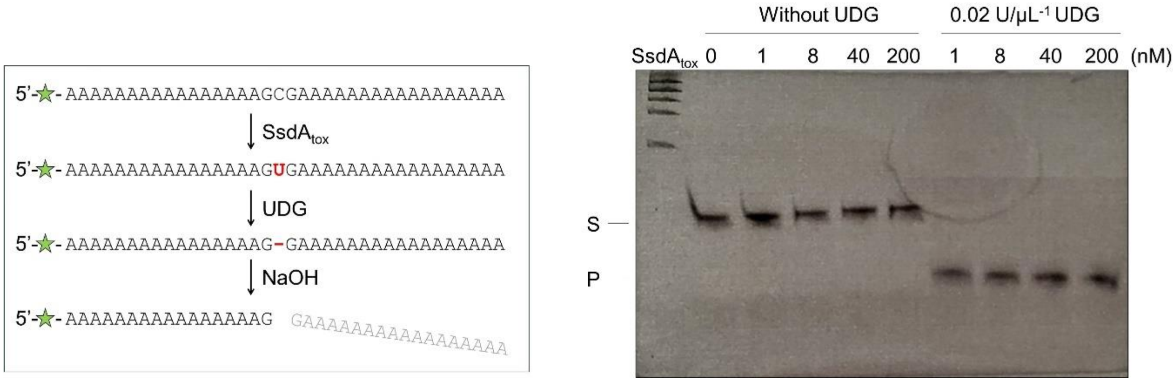


**Fig. S2. In vitro deaminase assay using SsdA$_{tox}$ protein.**

A 5'-FAM labeled ssDNA containing a single cytosine base nucleotide was used as the substrate for the SsdA$_{tox}$ protein. Cytosine deaminated by the SsdA$_{tox}$ domain is converted to uracil, which is then recognized and cleaved by UDG. NaOH treatment denatures the ssDNA at the cleaved position.

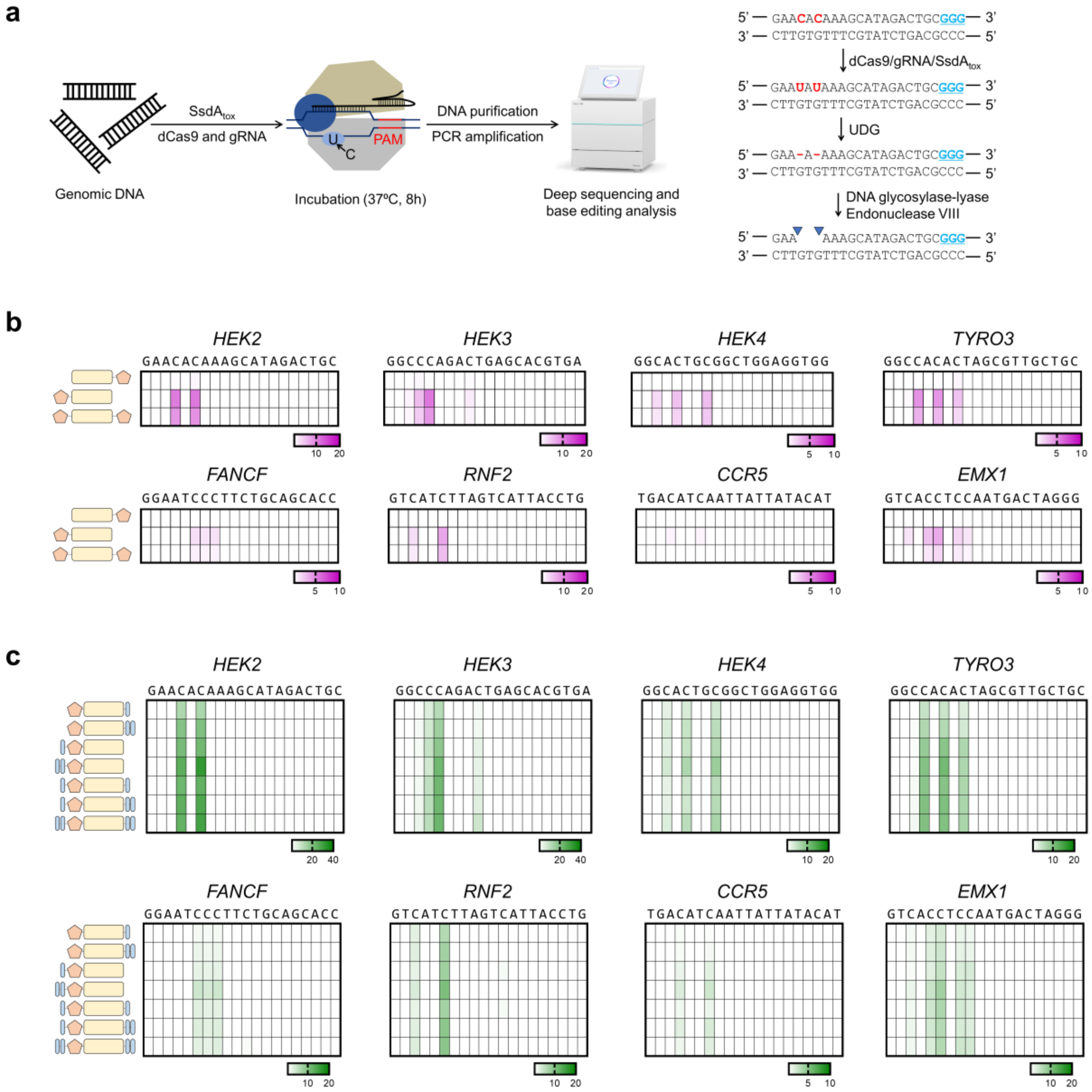


**Fig. S3. Developments of SsdAtox-based cytosine base editors.**

**a** Schematic overviews of in vitro deaminase assay. Genomic DNA from HEK293T/17 cells were subjected to in vitro deaminase assay and C-to-U conversion frequencies were measured by targeted-deep sequencing. **b, c** Base editing frequencies of each construct in HEK293T/17 *UNG* KO cell lines were described in heatmap. Data are presented as mean of independent biological triplicate (n=3).

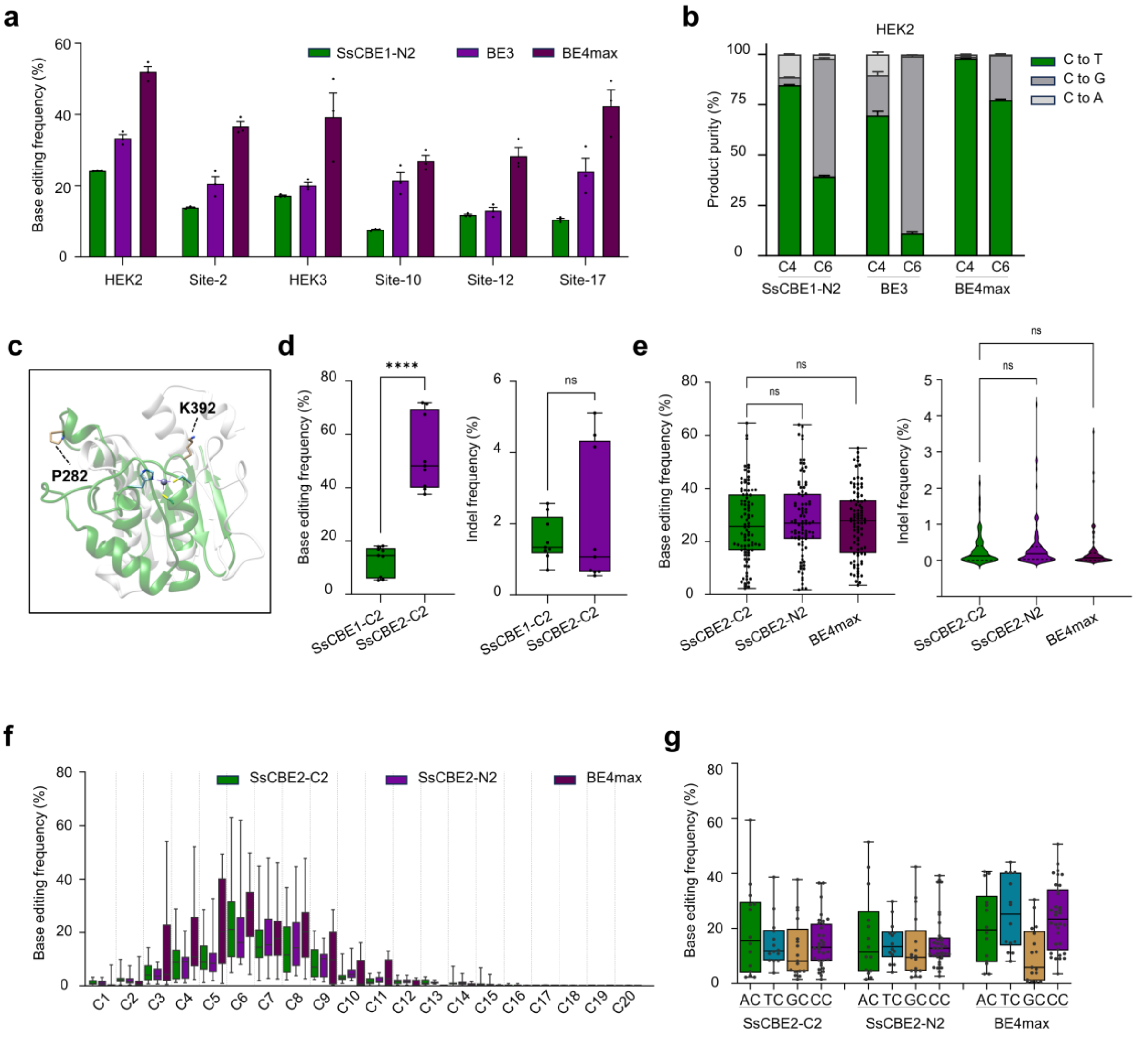


**Fig. S4. Rational engineering of $SsdA_{tox}$ domain.**

**a** Base editing frequency of SsCBE1-N2, BE3, and BE4max compared in HEK293T/17 cells. Data are presented as means of independent biological triplicate (n=3) and error bare means SEM. **b** Product purity comparison of SsCBE1-N2, BE3, and BE4max at HEK2 target site. **c** Structural alignments of TadA and $SsdA_{tox}$ domain highlighting two candidate residues for engineering: P282 and K392. **d** Comparison of base editing and indel frequency of SsCBE1-C2 and SsCBE2-UGI-C2 across three target sites in HEK293T/17 cells. **** $P<0.0001$ by unpaired *t*-test. **e** Comparison of base editing and indel frequency of SsCBE2-C2, SsCBE2-N2, and BE4max across 29 endogenous target sites in HEK293T/17 cells. Statistical analysis performed using unpaired *t*-test. **f, g** Base editing window and context analysis of three constructs across 29 endogenous target sites in HEK293T/17 cells.

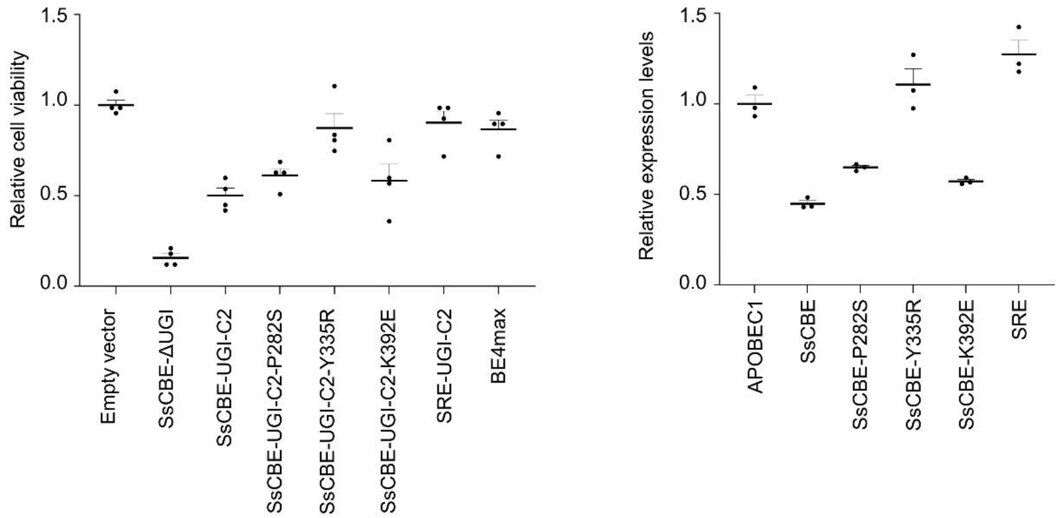


**Fig. S5. Cytotoxicity and expression of SsdA$_{tox}$ variants.**

To measure cell viability, each SsdA$_{tox}$ variant was transfected into HEK293T/17 cells, and the luminescent assay was conducted 72h after transfection. Relative cell viabilities were calculated by dividing the Relative Light Units (RLUs) of cells transfected with each variant by the RLUs of cells transfected with the empty vector. To compare expression level, the P2A-mehcrry fused each deaminase domain was transfected into HEK293T/17 cells and FACS analysis was performed. The relative expression levels were calculated by dividing the percentage of PerCP-Cy5-5-A positive cells transfected with each variant by the percentage of PerCP-Cy5-5-A positive cells transfected with the APOBEC1-P2A-mcherry construct. The transfection was conducted in independent biological triplicate (n=3).

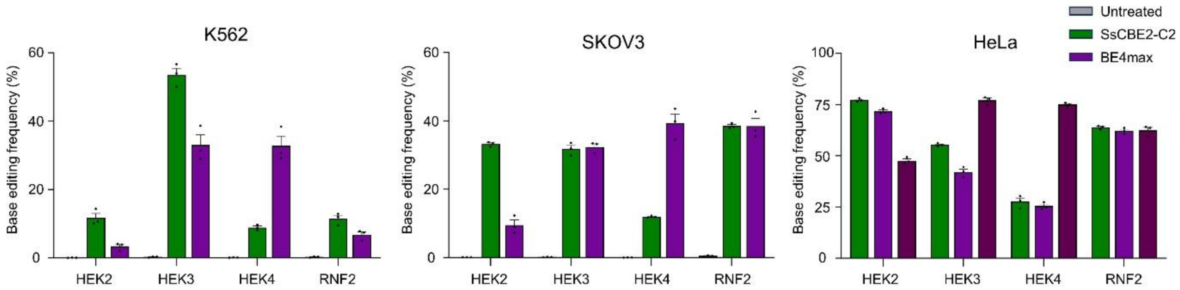


**Fig. S6. Base editing in other three cell lines.**

SsCBE2-C2 and BE4max were transfected in three different cell line and base editing frequencies were measured by targeted-deep sequencing across 4 target sites. Data are presented as mean and error bars means SEM of independent biological triplicate (n=3).

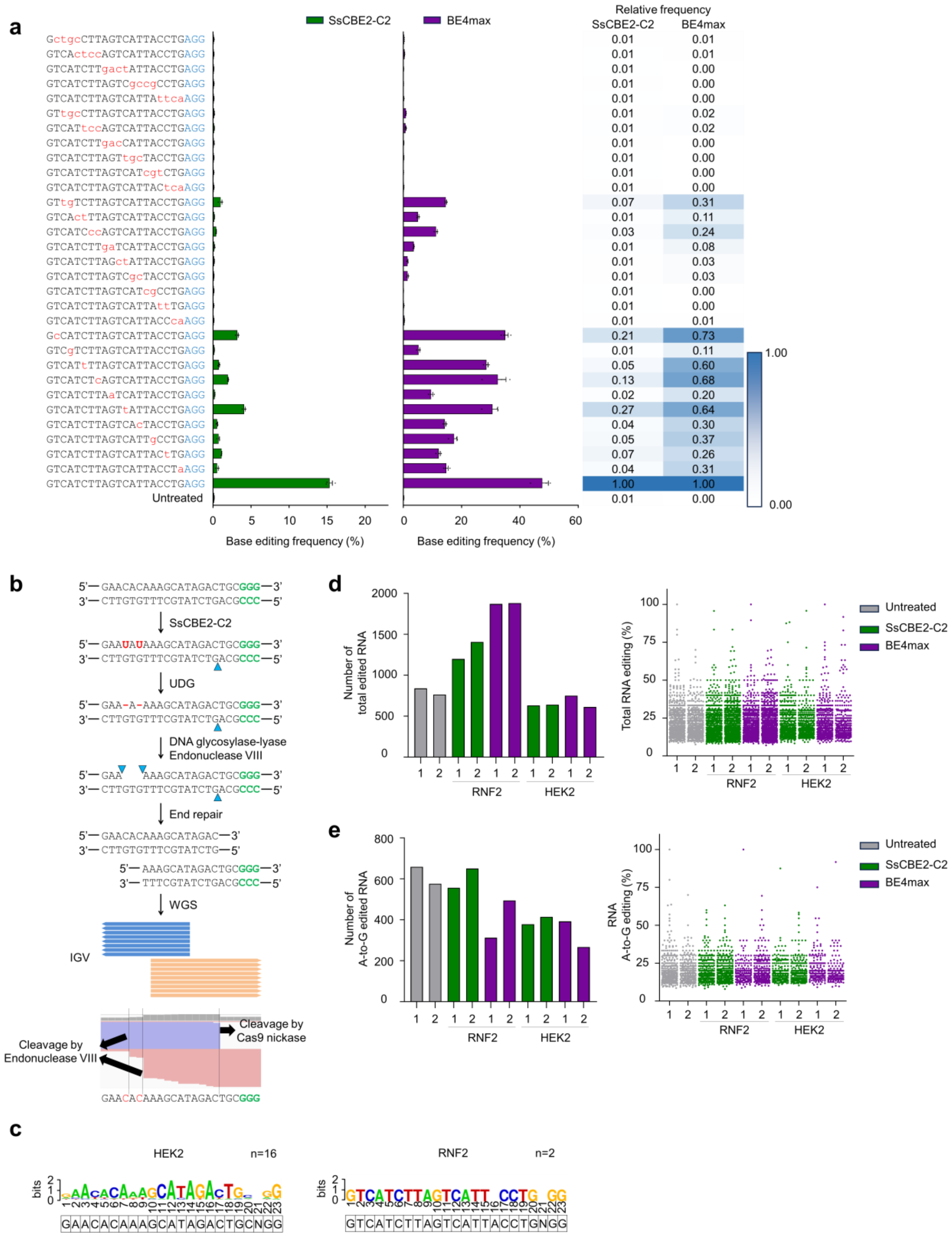
a
SsCBE2-C2
BE4max
Relative frequency
SsCBE2-C2 BE4max
GctgcCTTAGTCATTACCTGAGG 0.01 0.01
GTCActccAGTCATTACCTGAGG 0.01 0.01
GTCATCTTgactATTACCTGAGG 0.01 0.00
GTCATCTTAGTCgccgCCTGAGG 0.01 0.00
GTCATCTTAGTCATTAttcaAGG 0.01 0.00
GTtgcCTTAGTCATTACCTGAGG 0.01 0.02
GTCATtccAGTCATTACCTGAGG 0.01 0.02
GTCATCTTgacCATTACCTGAGG 0.01 0.00
GTCATCTTAGTtgcTACCTGAGG 0.01 0.00
GTCATCTTAGTCATcgtCTGAGG 0.01 0.00
GTCATCTTAGTCATTACtcaAGG 0.01 0.00
GTtgTCTTAGTCATTACCTGAGG 0.07 0.31
GTCActTTAGTCATTACCTGAGG 0.01 0.11
GTCATCccAGTCATTACCTGAGG 0.03 0.24
GTCATCTTgaTCATTACCTGAGG 0.01 0.08
GTCATCTTAGctATTACCTGAGG 0.01 0.03
GTCATCTTAGTCgcTACCTGAGG 0.01 0.03
GTCATCTTAGTCATcgCCTGAGG 0.01 0.00
GTCATCTTAGTCATTAttTGAGG 0.01 0.00
GTCATCTTAGTCATTACCcaAGG 0.01 0.01
GcCATCTTAGTCATTACCTGAGG 0.21 0.73
GTCgTCTTAGTCATTACCTGAGG 0.01 0.11
GTCATtTTAGTCATTACCTGAGG 0.05 0.60
GTCATCTcAGTCATTACCTGAGG 0.13 0.68
GTCATCTTAaTCATTACCTGAGG 0.02 0.20
GTCATCTTAGTtATTACCTGAGG 0.27 0.64
GTCATCTTAGTCAcTACCTGAGG 0.04 0.30
GTCATCTTAGTCATTgCCTGAGG 0.05 0.37
GTCATCTTAGTCATTACtTGAGG 0.07 0.26
GTCATCTTAGTCATTACCTaAGG 0.04 0.31
GTCATCTTAGTCATTACCTGAGG 1.00 1.00
Untreated 0.01 0.00
1.00
0.00
0 5 10 15 20
0 20 40 60
Base editing frequency (%)
Base editing frequency (%)
b
5'—GAACACAAAGCATAGACTGCGGG—3'
3'—CTTGTGTTTCGTATCTGACGCCC—5'
SsCBE2-C2
5'—GAAUAUAAAGCATAGACTGCGGG—3'
3'—CTTGTGTTTCGTATCTGACGCCC—5'
UDG
5'—GAA-A-AAAGCATAGACTGCGGG—3'
3'—CTTGTGTTTCGTATCTGACGCCC—5'
DNA glycosylase-lyase
Endonuclease VIII
5'—GAA AAAGCATAGACTGCGGG—3'
3'—CTTGTGTTTCGTATCTGACGCCC—5'
End repair
5'—GAACACAAAGCATAGAC—3'
3'—CTTGTGTTTCGTATCTG—5'
5'—AAAGCATAGACTGCGGG—3'
3'—TTTCGTATCTGACGCCC—5'
WGS
IGV
Cleavage by Cas9 nickase
Cleavage by Endonuclease VIII
GAACACAAAGCATAGACTGCGGG
c
HEK2 n=16
RNF2 n=2
bits
GAACACAAAGCATAGACTGCNGG
GTCATCTTAGTCATTACCTGNGG
d
Number of total edited RNA
2000
1500
1000
500
0
1 2 1 2 1 2 1 2 1 2
RNF2 HEK2
Total RNA editing (%)
100
75
50
25
0
Untreated
SsCBE2-C2
BE4max
e
Number of A-to-G edited RNA
800
600
400
200
0
RNA A-to-G editing (%)

**Fig. S7. Analysis of off-target effects of SsCBE2.**

**a** Tolerance evaluation of SsCBE2-C2 and BE4max for mismatched sgRNAs with one to four nucleotides mismatches from the RNF2 site in HEK293T/17 cells. PAM sequences are indicated in blue and mismatched bases are indicated in red. Relative frequencies were calculated by dividing base editing frequencies obtained with mismatched sgRNAs by the mean base editing frequency of the matched sgRNA. Data are presented as mean and error bar means SEM of independent biological triplicate (n=3). **b** Overviews of Digenome-seq using SsCBE2-C2 at HEK2 target site. SsCBE-C2 catalyzes C-to-U conversion and the uracil-containing sites were cleaved by USER enzyme, a mixture of *E.coli* uracil DNA glycosylase (UDG) and DNA glycosylase-lyase endonuclease VIII. Arrows indicate the positions of phosphodiester bonds cleaved by the SpCas9(D10A) nickase and USER. IGV image shows straight alignments of sequence reads at HEK2 on-target site. **c** Nucleotide sequences captured by Digenome-seq were compared and sequence logos were obtained using WebLogo. **d, e** Cas9-independent RNA off-target deamination of SsCBE2-C2 and BE4max in HEK293T/17 cells. Transcriptome sequencing was used to determine the number and frequency of total RNA editing, **d**, and of A-to-G RNA editing, **e**.

a

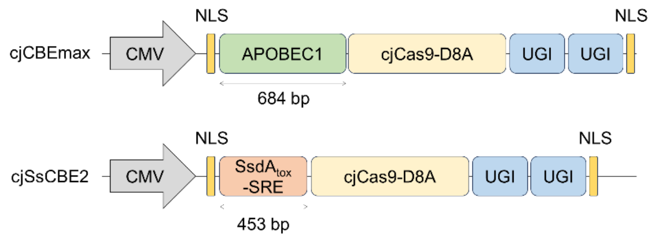


b

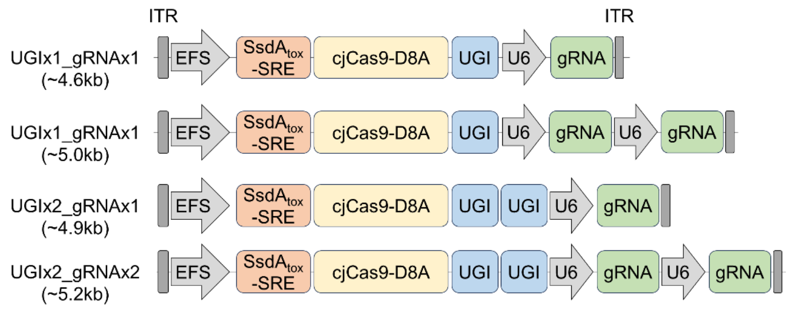


**Fig. S8. Architectures of cjCas9-based cytosine base editors.**

**a** Schematic overview of cjSsCBE2 and cjCBEmax. **b** Schematic overviews of single AAV vectors encoding various cjSsCBE2 variants and gRNA. The length between ITR is described in each construct.

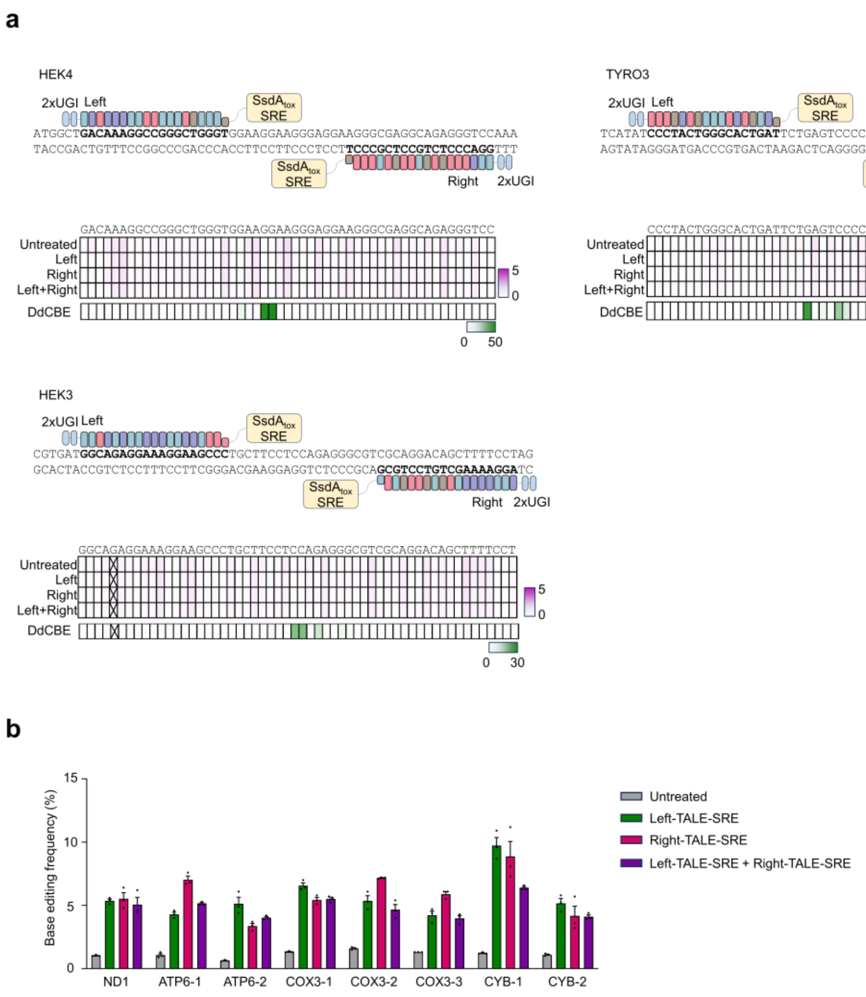


**Fig. S9. Base editing using TALE-SRE.**

**a** Base editing frequency of TALE-SRE targeting nucleus genome. Monomer or dimer forms of TALE-SRE were transfected in HEK293T/17 cells and DdCBE was used as positive controls. Transfection was conducted in biological triplicate and base editing frequencies were described in heatmaps. **b** Base editing frequencies by TALE-SRE in mitochondrial genome. Data are presented as mean and error bars mean SEM of independent biological triplicate (n=3).

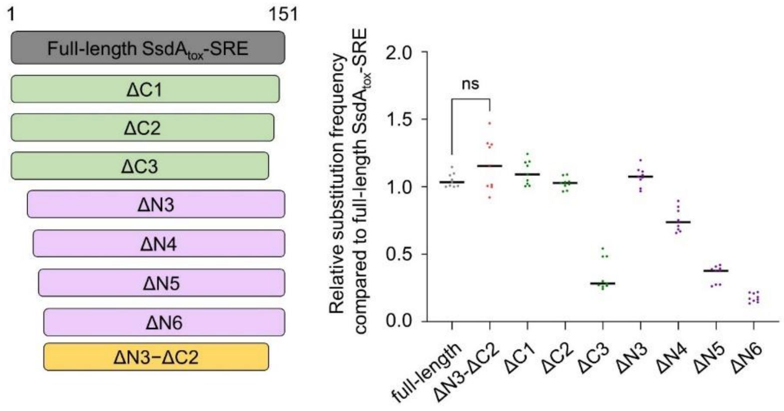


**Fig. S10. Truncation of SsdA$_{tox}$-SRE domain.**

Either N-terminally or C-terminally truncated SsdA$_{tox}$-SRE domains were cloned into SsCBE2-C2, and their base editing frequencies across three target sites in HEK293T/17 cells were measured by targeted-deep sequencing. Relative frequencies were calculated by dividing base editing frequencies obtained with each variant by the mean base editing frequency obtained with full-length SsdA$_{tox}$-SRE in each target site.

### Table S1. Identified potential off-target sites by Digenome-seq.

| HEK2 | Chr. | Location | DNA cleavage score | DNA seq at a cleavage site | Bulge |
|---|---|---|---|---|---|
| HEK2_OT1 | chr.5 | 87240613 | 14.2 | GAACACAAAGCATAGACTGCGGG | X |
| HEK2_OT2 | chr.15 | 93557679 | 7.5 | GAACACA-tGCATAGACTGCTAG | O |
| HEK2_OT3 | chr.4 | 90522183 | 6.7 | GAACACAAtGCATAGAtTGCCGG | X |
| HEK2_OT4 | chr.2 | 19844956 | 1.3 | aActcCAAAGCATAtACTGCTGG | X |
| HEK2_OT5 | chr.13 | 55564918 | 1.2 | acACACAAAGCAT-GACTGCAGG | X |
| HEK2_OT6 | chr.1 | 167742859 | 0.7 | aAACACAgAGCAcAGACTGCTGA | X |
| HEK2_OT7 | chr.19 | 35505485 | 0.6 | GAACAC-AAGCAcAGACTGaAGG | O |
| HEK2_OT8 | chr.1 | 36097072 | 0.4 | GtAaACAAAGCATAGACTGaGGG | X |
| HEK2_OT9 | chr.2 | 192248363 | 0.4 | GAACACAtA-CATAGACaGCTGG | X |
| HEK2_OT10 | chr.11 | 128508576 | 0.3 | GAAttCAAAGCATAGAtTGCAGG | X |
| HEK2_OT11 | chr.1 | 77190607 | 0.3 | tcACACAAAcCATAGACTGaGGG | X |
| HEK2_OT12 | chr.4 | 135329594 | 0.3 | tAACAaAtAGCATAGACTGtGTG | X |
| HEK2_OT13 | chr.8 | 97317606 | 0.3 | GAACACAgtaCATAGACTGgCAG | X |
| HEK2_OT14 | chr.9 | 290167 | 0.2 | aAACAtAAAGaATAGACTGCAAG | X |
| HEK2_OT15 | chr.4 | 53536209 | 0.2 | GAAtACtAAGCATAGACTcCAGG | X |
| HEK2_OT16 | chr.19 | 28824655 | 0.2 | GAACtCAAAGCATAGAaTaaTGG | X |

| RNF2 | | | | | |
|---|---|---|---|---|---|
| RNF2_OT1 | chr.1 | 185056773 | 3.1 | GTCATCTTAGTCATTACCTGAGG | X |
| RNF2_OT2 | chr.10 | 75832488 | 0.5 | GtCATCTTAGTCATT-CCTGGGG | O |

### Table S2. Nucleotide sequences of target sites and PCR primers.

| Target sites | Spacer sequences | PAM | PCR-F | PCR-R |
|---|---|---|---|---|
| AAVS1 | GCTGACTCAGAGACCCTGAG | TGG | GGCCCAGACTAGCCCAGTTGT | CCACCTGCCTTGGCCTCTCA |
| CCR5 | TGACATCAATTATTATACAT | CGG | GAGGGCAACTAAATACATTCTAGGAC | CCAAAGATGAACACCAGTGA |
| CUL3 | GTAAACCTGGAATAACACGA | TGG | TTGGGAGCACTTCCAGGTTCACT | CTGCACTCCAGCCTTGGTGACAG |
| EMX1 | GTCACCTCCAATGACTAGGG | TGG | GGACAAAGTACAAACGGCAGA | AGTGGCCAGAGTCCAGCTT |
| FANCF | GGAATCCCTTCTGCAGCACC | TGG | ATGGATGTGGCGCAGGTAG | AGCATTGCAGAGAGGCGTAT |
| HEK2 | GAACACAAAGCATAGACTGC | GGG | AGACCTGGCTGAGCTAACTG | TCCAGCCCCATCTGTCAAAC |
| HEK3 | GGCCCAGACTGAGCACGTGA | TGG | GCATGCATTTGTAGGCTTGA | CCCAGCCAAACTTGTCAAC |
| HEK4 | GGCACTGCGGCTGGAGGTGG | GGG | CTCCCTTCAAGATGGCTGAC | AACGGAGACACACACACAGG |
| RNF2 | GTCATCTTAGTCATTACCTG | AGG | ATTTCCAGCAATGTCTCAGG | GCCAACATACAGAAGTCAGGAA |
| TYRO3 | GGCCACACTAGCGTTGCTGC | TGG | TCCCTACTGGGCACTGATTC | TCCCTGTCAACAAAGTGCTG |
| Site-1 | CCAGCCCGCTGGCCCTGTAA | AGG | AGACCTGGCTGAGCTAACTG | TCCAGCCCCATCTGTCAAAC |
| Site-2 | GCTGGCCCTGTAAAGGAAAC | TGG | AGACCTGGCTGAGCTAACTG | TCCAGCCCCATCTGTCAAAC |
| Site-3 | GTTTCCTTTACAGGGCCAGC | GGG | AGACCTGGCTGAGCTAACTG | TCCAGCCCCATCTGTCAAAC |
| Site-4 | GCACTTGTTTGCAGCTATTC | AGG | AGACCTGGCTGAGCTAACTG | TCCAGCCCCATCTGTCAAAC |
| Site-5 | CTGCTTCTCCAGCCCTGGCC | TGG | GCATGCATTTGTAGGCTTGA | CCCAGCCAAACTTGTCAAC |
| Site-6 | CCCTGGCCTGGGTCAATCCT | TGG | GCATGCATTTGTAGGCTTGA | CCCAGCCAAACTTGTCAAC |
| Site-7 | GGAAGCCCTGCTTCCTCCAG | AGG | GCATGCATTTGTAGGCTTGA | CCCAGCCAAACTTGTCAAC |
| Site-8 | CTTCCTCCAGAGGGCGTCGC | AGG | GCATGCATTTGTAGGCTTGA | CCCAGCCAAACTTGTCAAC |
| Site-9 | CAGGACAGCTTTTCCTAGAC | AGG | GCATGCATTTGTAGGCTTGA | CCCAGCCAAACTTGTCAAC |
| Site-10 | CAGCTCCTGCACCGGGATAC | TGG | GCATGCATTTGTAGGCTTGA | CCCAGCCAAACTTGTCAAC |
| Site-11 | GGGGCACCGCGGCGCCCCGG | TGG | CTCCCTTCAAGATGGCTGAC | AACGGAGACACACACACAGG |
| Site-12 | GCGGCGCCCCGGTGGCACTG | CGG | CTCCCTTCAAGATGGCTGAC | AACGGAGACACACACACAGG |
| Site-13 | CGCCCCGGTGGCACTGCGGC | TGG | CTCCCTTCAAGATGGCTGAC | AACGGAGACACACACACAGG |
| Site-14 | TCCCTTCCTTCCACCCAGCC | CGG | CTCCCTTCAAGATGGCTGAC | AACGGAGACACACACACAGG |
| Site-15 | CCCTGCCTGTCATCCTGCTT | TGG | CTCCCTTCAAGATGGCTGAC | AACGGAGACACACACACAGG |
| Site-16 | GCAGTGCCACCGGGGCGCCG | CGG | CTCCCTTCAAGATGGCTGAC | AACGGAGACACACACACAGG |
| Site-17 | CTCCAGCCGCAGTGCCACCG | GGG | CTCCCTTCAAGATGGCTGAC | AACGGAGACACACACACAGG |
| Site-18 | ACCTCCAGCCGCAGTGCCAC | CGG | CTCCCTTCAAGATGGCTGAC | AACGGAGACACACACACAGG |
| Site-19 | TACACGTCTCATATGCCCCT | TGG | ATTTCCAGCAATGTCTCAGG | GCCAACATACAGAAGTCAGGAA |
| Site-20 | TCAACCATTAAGCAAAACAT | GGG | ATTTCCAGCAATGTCTCAGG | GCCAACATACAGAAGTCAGGAA |
| Rloop-site-1 | GTGGTAGACAGCATGTGTCCTA | AAGGG | TCCTGCAGTCTCCTGCTTCT | ACCAACATACATGCCCCTTT |
| Rloop-site-2 | ATTTACAGCCTGGCCTTTGGGG | TCGGG | GACATTTCCACCGCAAAATG | CGGTGGGAGATCTGGTTTC |
| Rloop-site-3 | GTGTCAGGTAATGTGCTAAACA | GAGAG | TGCTCCAGATTTCCCTTCAT | GGCATCCAGAGACATGGTTT |
| Rloop-site-4 | TCTGCTTCTCCAGCCCTGGC | CTGGG | AAACGCCCATGCAATTAGTC | CAGGAGCTGCACATACTAGCC |
| Rloop-site-5 | GATGTTCCAATCAGTACGCA | GAGAG | GAAAAGCGATCCAGGTGCT | GGCTTTTAAGTTGCCCAGAG |
| EPAS | CAGGACAGCAGGGGCTCCTTGT | AGCCACAC | AAGCCTTGGAGGGTTTCATT | GTGGCTAGCACCTTCCACTC |
| HIF | GCTATTCACCAAAGTTGAATCA | GAAGATAC | CCCTTCCCCTCACTGTATCA | GGCCAGCAAAGTTAAAGCAT |
| ANGPT2 | GCTGTGCAGAGGGACGCGCCGC | TCGAATAC | ATGGGTCCTGCAGCTACACT | TTCCATGATGTTCTCCAGCA |
| HPD-1 | TTTTCCACCCGTAGTATGGGGA | CACCACAC | GGAAGTAGGGGGTCCATGA | ACGCATCTGGTTAGGGTCAG |
| HPD-2 | TTCCACCCGTAGTATGGGGACA | CCACACAC | GGAAGTAGGGGGTCCATGA | ACGCATCTGGTTAGGGTCAG |
| mTyr | ACCTCAGTTCCCCTTCAAAG | GGG | AACCCATGAAGTTGCCTGAG | TTGTTGGCAAAAGAATGCTG |

### Table S3. Target sequences and PCR sequences for TALE-SRE.

| Target sites | Target sequence | PCR-F | PCR-R |
|---|---|---|---|
| ND1-Left | AGCCGTTTACTCAATCCTC | GCTCTCACCATCGCTCTTCT | TGATGGCTAGGGTGACTTCAT |
| ND1-Right | CAGGGCGTAGTTTGAGTTT | GCTCTCACCATCGCTCTTCT | TGATGGCTAGGGTGACTTCAT |
| ATP6-1-Left | CAACAACCGACTAATCACC | CCCTCTATTGATCCCCACCT | GATGGCCATGGCTAGGTTTA |
| ATP6-2-Right | TTGTTTTGAGGTTAGTTTG | CCCTCTATTGATCCCCACCT | GATGGCCATGGCTAGGTTTA |
| ATP6-2-Left | AACCATACACAACACTAAA | CCCTCTATTGATCCCCACCT | GATGGCCATGGCTAGGTTTA |
| ATP6-2-Right | GATTAAGGATACTAGTATA | CCCTCTATTGATCCCCACCT | GATGGCCATGGCTAGGTTTA |
| COX3-1-Left | ATCATATAGTAAAACCCAG | CAACACATAATGACCCACCAA | GAAGGCCTTTTTGGACAGGT |
| COX3-1-Right | AGGAGGGCTGAGAGGGCCC | CAACACATAATGACCCACCAA | GAAGGCCTTTTTGGACAGGT |
| COX3-2-Left | CCTAATGACCTCCGGCCTA | CAACACATAATGACCCACCAA | GAAGGCCTTTTTGGACAGGT |
| COX3-2-Right | GAGGAGCGTTATGGAGTGG | CAACACATAATGACCCACCAA | GAAGGCCTTTTTGGACAGGT |
| COX3-3-Left | AACCAACACACTAACCATA | CAACACATAATGACCCACCAA | GAAGGCCTTTTTGGACAGGT |
| COX3-3-Right | ATGTGCTTTCTCGTGTTAC | CAACACATAATGACCCACCAA | GAAGGCCTTTTTGGACAGGT |
| CYB-1-Left | CCAACATCTCCGCAT | AACCACTCATTCATCGACCTC | CGCCCGATGTGTAGGAAG |
| CYB-1-Right | CAGGCAGGCGCCAAGGAGT | AACCACTCATTCATCGACCTC | CGCCCGATGTGTAGGAAG |
| CYB-2-Left | CGAGACGTAAATTATGGC | AACCACTCATTCATCGACCTC | CGCCCGATGTGTAGGAAG |
| CYB-2-Right | GAGGCGCCATTGGCGTGAA | AACCACTCATTCATCGACCTC | CGCCCGATGTGTAGGAAG |
| HEK4-Left | GACAAAGGCCGGGCTGGGT | CTCCCTTCAAGATGGCTGAC | AACGGAGACACACACACAGG |
| HEK4-Right | GGACCCTCTGCCTCGCCCT | CTCCCTTCAAGATGGCTGAC | AACGGAGACACACACACAGG |
| TYRO3-Left | CCCTACTGGGCACTGAT | TCCCTACTGGGCACTGATTC | TCCCTGTCAACAAAGTGCTG |
| TYRO3-Right | GAAGGGGGCTGCAGGCAGT | TCCCTACTGGGCACTGATTC | TCCCTGTCAACAAAGTGCTG |
| HEK3-Left | GGCAGAGGAAAGGAAGCCC | GCATGCATTTGTAGGCTTGA | CCCAGCCAAACTTGTCAAC |
| HEK3-Right | AGGAAAAGCTGTCCTGCG | GCATGCATTTGTAGGCTTGA | CCCAGCCAAACTTGTCAAC |
